# CLN7 mutation causes aberrant redistribution of protein isoforms and contributes to Batten disease pathobiology

**DOI:** 10.1101/2022.04.21.488782

**Authors:** Aseel M. Sharaireh, Marta Guevara-Ferrer, Saul Herranz-Martin, Marina Garcia-Macia, Alexander Phillips, Anna Tierney, Michael P Hughes, Oliver Coombe-Tennant, Hemanth Nelvagel, Alysha E. Burrows, Stuart Fielding, Lorna M. FitzPatrick, Christopher D. Thornton, Stephan Storch, Sara E. Mole, Andrew Dowsey, Richard Unwin, Juan P. Bolanos, Ahad A. Rahim, Tristan R. McKay

## Abstract

The variant late infantile form of the inherited neurodegenerative Batten disease (BD) is caused by mutations in the CLN7/MFSD8 gene and represents a strong candidate for gene therapy. Post-natal intracerebral administration of AAV9-hCLN7 to *Cln7*^Δex2^ knockout mice resulted in extended lifespan but dose escalation resulted in reduced acuity in neurophysiology tests, cerebral atrophy and elevated neuroinflammation. Comparing patient and control iPSC-derived neural progenitor cells (iNPC) we discovered that CLN7 localizes to the nucleus as well as the endolysosomal network and is differentially distributed in BD iNPC. Proteomics identified a profound nuclear defect in BD iNPC that compounds with mitochondrial and lysosomal metabolic defects resulting in elevated apoptosis. We further identified a 50kDa common nuclear CLN7 isoform and a 37kDa isoform that accumulates only in BD iNPC nuclei. Our findings suggest that successful treatment of CLN7 BD will require combinatorial therapies addressing both loss and aberrant gain of protein function.

## Introduction

The Neuronal Ceroid Lipofuscinoses (NCLs), also known as Batten disease (BD), are a group of inherited autosomal recessive monogenic neurodegenerative disorders caused by mutations in *CLN* genes with a collective incidence of up to 1:12,500 live births. Clinical presentation is almost exclusively in the first decade of life with common clinical features that include seizures, visual failure, and cognitive and motor decline due to progressive cerebral atrophy. The NCLs are sub-categorized by age-of-onset into infantile, late infantile, juvenile and rare adult-onset forms with a shared neuropathology of lysosomal lipofuscin accumulation in cortical neurons preceding neurodegeneration (*1*, *2*). Most CLN proteins localize to vesicular transport and autophagic/lysosomal degradation networks, but many remain functionally undefined. At least four are soluble lysosomal enzymes (CLN1/PPT1, CLN2/TPP1, CLN5, CLN10/CTSD) and Cerliponase alfa has proven to be a safe and efficacious enzyme replacement therapy in CLN2 Phase I/II clinical trials (*3*). CLN6 and CLN8 are ER-Golgi lysosomal enzyme vesicular transport transmembrane proteins (*4*, *5*) CLN3 and CLN7 (also known as MFSD8) are transmembrane proteins belonging to the MFS superfamily (*6*) and found in the lysosome perimeter membrane, with CLN7 recently shown to be a chloride channel mediating the movement of chloride ions from the lysosome into the cytosol (*7*). These membrane proteins represent good potential candidates for gene therapy. Pre-clinical studies applying AAV vectors expressing full-length cDNAs to the central nervous system (CNS) by various routes of administration have shown indicators of efficacy (*8*–*12*), leading the way to clinical trial approvals for CLN3, CLN6 and CLN7. Most patients with CLN3 disease share a common 1kb intragenic deletion (*13*) and alternative splicing has been exploited using antisense oligonucleotide (ASO)-mediated exon skipping to engineer a partially functional CLN3 isoform capable of ameliorating some aspects of disease in patient cells and the *Cln3*^Δex7/8^ mouse model (*14*). Furthermore, modelling of the predominant *CLN3*^Δex7/8^ 1-kb intragenic deletion in the yeast orthologue *Btn1* implies multiple gain-of-function activities for truncated protein isoforms generated by alternative splicing (*15*) implying that there are both positive and negative consequences of splice isoforms in NCL. Interestingly, like CLN7, CLN3 can also be considered a member of the MFS family (*6*). Improved understanding of molecular pathogenicity has enabled the identification of tamoxifen (*16*) and the PFKFB3 inhibitor AZ67 (*17*) as FDA approved repurposed drugs capable of ameliorating aspects of CLN7 BD in patient iNPCs and *Cln7^Δex2^* mice. Overall, a complete understanding from gene expression to protein functions will facilitate targeted and efficient therapies for the NCLs.

CLN7 disease presents predominantly in late infancy, and up to adulthood. There are more than 50 underlying mutations with most causing severe disease presumed to be due to loss of function. Others are associated with juvenile or later ages of onset or a significantly milder disease course dominated by retinal dystrophy (*13*). CLN7 disease presents as a strong candidate for brain-directed gene therapy (*1*). In our own investigations we have found that neonatal intracranial administration of an AAV9 vector expressing the full length hCLN7 using the strong synapsin promoter was only able to partially correct CNS disease in the well-characterized *Cln7^Δex2^* mouse model (*18*) with the emergence of dose-dependent neuroinflammation and astrogliosis. This unexpected observation in these gene therapy studies led us to conduct a comprehensive evaluation of disease phenotype in iNPC from two CLN7 disease patients presenting in late infancy, reported here, which revealed CLN7 has multiple functionally distinct isoforms that are differentially localized in BD cells compared to control iNPC from age-matched individuals. CLN7 localizes to the autophagic/lysosomal vesicular network and nucleus but dynamically redistributes in an opposing manner between normal and BD iNPC following treatment with the vATPase inhibitor bafilomycin A (Baf A). This is concomitant with mitochondrial defects including elevated reactive oxygen species (ROS), deficits in oxidative phosphorylation (OXPHOS) and elevated cytochrome c-mediated apoptosis. A comprehensive proteomic evaluation confirms a BD stress response triangulating mitochondrial bioenergetics, vesicular transport, and a profound nuclear defect with a pronounced depletion of nuclear pore complex proteins. A 50kDa CLN7 isoform (CLN7^50kDa^) localizes to the nucleus in control iNPCs but there is a further 37kDa isoform (CLN7^37kDa^) localizing to the nucleus in BD iNPCs. We have extrapolated pathological phenotypes common to the *Cln7^Δex2^* mouse and CLN7 iNPC attributed to a CLN7 loss-of-function and those unique to CLN7 iNPC that are more likely caused by malevolent gain-of-function. Moreover, we have identified distinct normal and disease-associated nuclear isoforms of CLN7 that strongly imply a novel nuclear function that is disrupted in BD and thus relevant to the development of effective treatments.

## Results

### AAV9-mediated gene therapy enhances survival of *Cln7^Δex2^* mice

Towards brain-directed gene therapy for CLN7 disease we conducted a comprehensive pre-clinical dose escalation, intracranial AAV9 gene therapy in neonatal mice. Neonatal (P0-P1) *Cln7^Δex2^* mouse pups received either a lower-dose (LD, 5μl of 2.5×10^10^vg n=18) or higher-dose (HD, 2.5×10^11^vg, n=11) of AAV9.SYN1.hCLN7 (AAV-CLN7) or 2.5×10^10^ vg of AAV9.SYN1.eGFP (AAV-GFP; n=11) via bilateral intracerebroventricular injections. Age-matched wild-type mice (n=22) and untreated *Cln7^Δex2^* mice (n=27) acted as controls. The humane endpoint was defined as when mice had lost 15% of body weight or recorded a poor neurological welfare score (*19*). *Cln7^Δex2^* mice treated with AAV-CLN7 injections demonstrated significantly increased lifespan when compared to untreated *Cln7^Δex2^* mice. However, while some HD-treated *C1n7^Δex2^* mice lived longer than the LD-treated *Cln7^Δex2^* mice, the median lifespan of both groups was not significantly different (**Fig. 1A**). Mice injected with control AAV-GFP vector showed no change in survival as compared to untreated *Cln7^Δex2^* mice. There was also no significant difference in body weight between any of the groups at any of the timepoints measured (**Supp. Fig. 2**).

**Figure 1.**
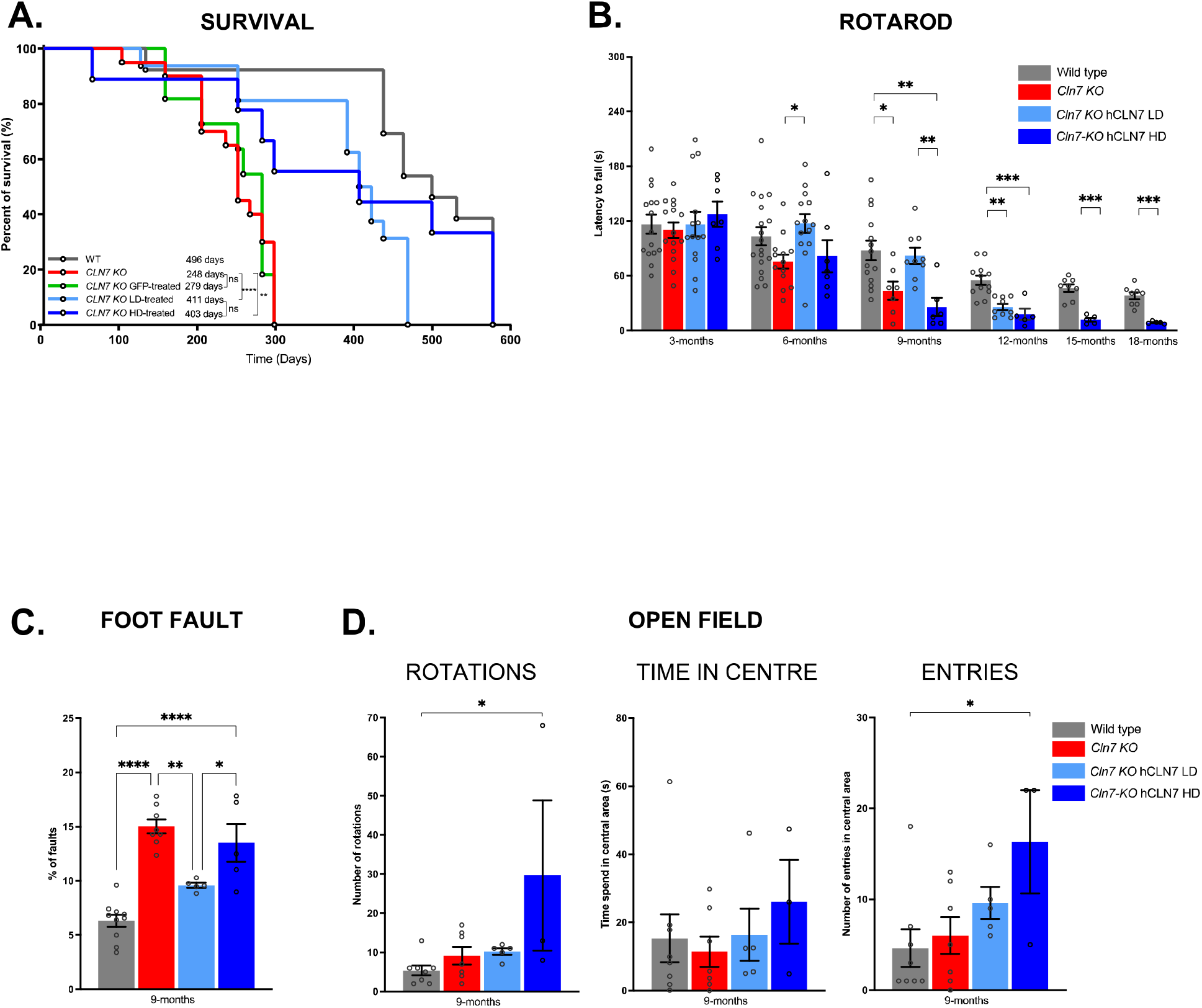
Dosage-dependent therapeutic effects of neonatal intracerebroventricular AAV9-hCLN7 gene therapy in *Cln7’^ex2^* mice. **(A)** Kaplan–Meier survival curve showing percentage of survival over time with median survival values for wild-type (WT), untreated *Cln7^Δex2^* mice (CLN7 KO), AAV9.SYN1.eGFP-treated *Cln7^Δe2^* mice (GFP), low-dose (LD) and high-dose (HD) AAV9.SYN1.hCLN7 treated *Cln7^Δex2^* mice (CLN7 KO) analyzed using Log-rank test (Mantel–Cox). **(B)** Accelerating rotarod performance measuring latency to fall, measured at 3-month intervals analyzed using two-way ANOVA with Tukey post-hoc correction. **(C)** Foot-fault measured as percentage of misplaced steps at 9 months analyzed using a one-way ANOVA with Tukey post-hoc correction. **(D)** Open-field behavioural analyses 9-months post-transduction quantified over 4 minutes by measuring the number of rotations and time spent in central area(s) and number of entries into central area analyzed using a one-way ANOVA with Tukey post-hoc correction.

### Gene therapy ameliorates behavioral deficits but efficacy is impeded at high-dose

Behavioral assessment of motor function demonstrated that some *Cln7^Δex2^* mice treated with AAV-CLN7 showed increased latency to fall on the accelerating rotarod paradigm from 6-9 months compared to *Cln7^Δex2^* mice without treatment who also rapidly increased in mortality. Interestingly, LD-treated mice significantly out-performed HD AAV-CLN7 treated *Cln7^Δex2^* mice during this period, being indistinguishable from wild-types (**Fig. 1B, Supp. Fig. 3A**). Similarly, foot fault assessment showed significant improvement in *Cln7^Δex2^* mice treated with LD but not in HD AAV-CLN7 (**Fig. 1C, Supp. Fig. 3B**). Our neurophysiology evaluations show some indications of gene therapy efficacy up to a decline between 9-12 months. Focusing our comparisons at 9-months, consistent with peak therapeutic efficacy and *Cln7^Δex2^* mouse decline, LD AAV-CLN7 showed increased therapeutic efficacy when compared to HD in *Cln7^Δex2^* mice by rotarod (**Fig. 1B)**, foot fault (**Fig. 1C)** and open field assessments (**Fig. 1D**). Both treatment groups showed improved performance on the vertical pole test as compared to untreated *Cln7^Δex2^* mice until 9 months, but thereafter showed decreased performance as compared to wild-type (**Supp. Fig. 3C**). Collectively, these analyses indicate that despite improved survival, AAV-CLN7 treated mice showed only moderate improvements in behavioral tests. Surprisingly, HD-treated mice were consistently outperformed by their LD-treated counterparts and under some criteria performed worse than untreated *Cln7^Δex2^* mice suggesting dose-related limitations to a gene therapy in this mouse model when expressing high levels of CLN7 using a strong promoter.

### Gene therapy ameliorates neuropathology but causes cortical thinning at high doses

Mice were sacrificed at 9 or 14-months and brain sections analyzed for cortical thickness as well as transgene expression and indicators of therapeutic efficacy or toxicity in the somatosensory barrel field of the cortex (S1BF). Cortical thickness was quantified using NeuN immunohistochemistry (IHC) and automated quantitation. At 9-months there was a trend toward increased cortical thickness at the S1BF in LD treated *Cln7^Δex2^* mice but a pronounced decrease in cortical thickness in HD treated mice beyond the underlying decrease of untreated mice suggestive of AAV-CLN7 dose-related neurodegeneration (**Fig. 2A**). Endogenous Cln7 expression was widespread in S1BF of wild-type mouse brains at 9-months with punctate immunolocalization that partially overlapped with lysosome-associated membrane protein 2 LAMP2. This is consistent with previously CLN7/MFSD8 immunolocalization detected throughout the vesicular endolysosomal network and nucleus (*7*, *20*). Massive over-expression of CLN7 in AAV-CLN7 transduced wild-type cells was exclusively co-localized with LAMP2 suggesting that transgenic CLN7 protein is restricted to the lysosome, in contrast to endogenous Cln7 (**Fig. 2B**). Additionally, there was clear evidence of AAV-CLN7 dose escalation translating to elevated levels of brain transduction and CLN7 expression in S1BF in *Cln7^Δex2^* mice (**Supp. Fig. 4A**). As expected, lysosome-associated membrane protein 1 (LAMP1) IHC was significantly higher in *Cln7^Δex2^* mice compared to wild-type, possibly due to the accumulation of defective lysosomes. However, although not statistically significant overall, LAMP1 IHC quantity decreased with LD but significantly increased with HD AAV-CLN7 treatment (**Supp. Fig. 4A**). By 14-months there was similarly high levels of LAMP1 immunostaining in both LD and HD treated *Cln7^Δex2^* mice (**Supp. Fig. 4B**). The accumulation of autofluorescent storage material (AFSM) is a well-established marker of lysosomal lipofuscin accumulation in cortical neurons affected by BD (*18*). AFSM was detectable at 9-months in S1BF *Cln7^Δex2^* mouse neurons compared to wild-type and both AAV-CLN7 treatment groups similarly reduced AFSM without reaching statistical significance. Similarly, we observed a significant increase in markers of astrocyte (GFAP) and microglial (CD68) activation in *Cln7^Δex2^* mice compared to wild-type. AAV-CLN7 gene therapy at LD ameliorated both markers of neuroinflammation but HD did not (**Fig. 2C & D**). At 14-months, where AAV-CLN7 treated *Cln7^Δex2^* mice survive but untreated mice do not, the levels of neuroinflammation are similarly high in both LD and HD treated mice (**Supp. Fig. 5**). Collectively, these observations present indications of a more efficacious gene therapy with LD AAV-CLN7 compared to HD in the *Cln7^Δex2^* mouse at 9-months that wanes at 14-months despite increasing lifespan.

**Figure 2.**
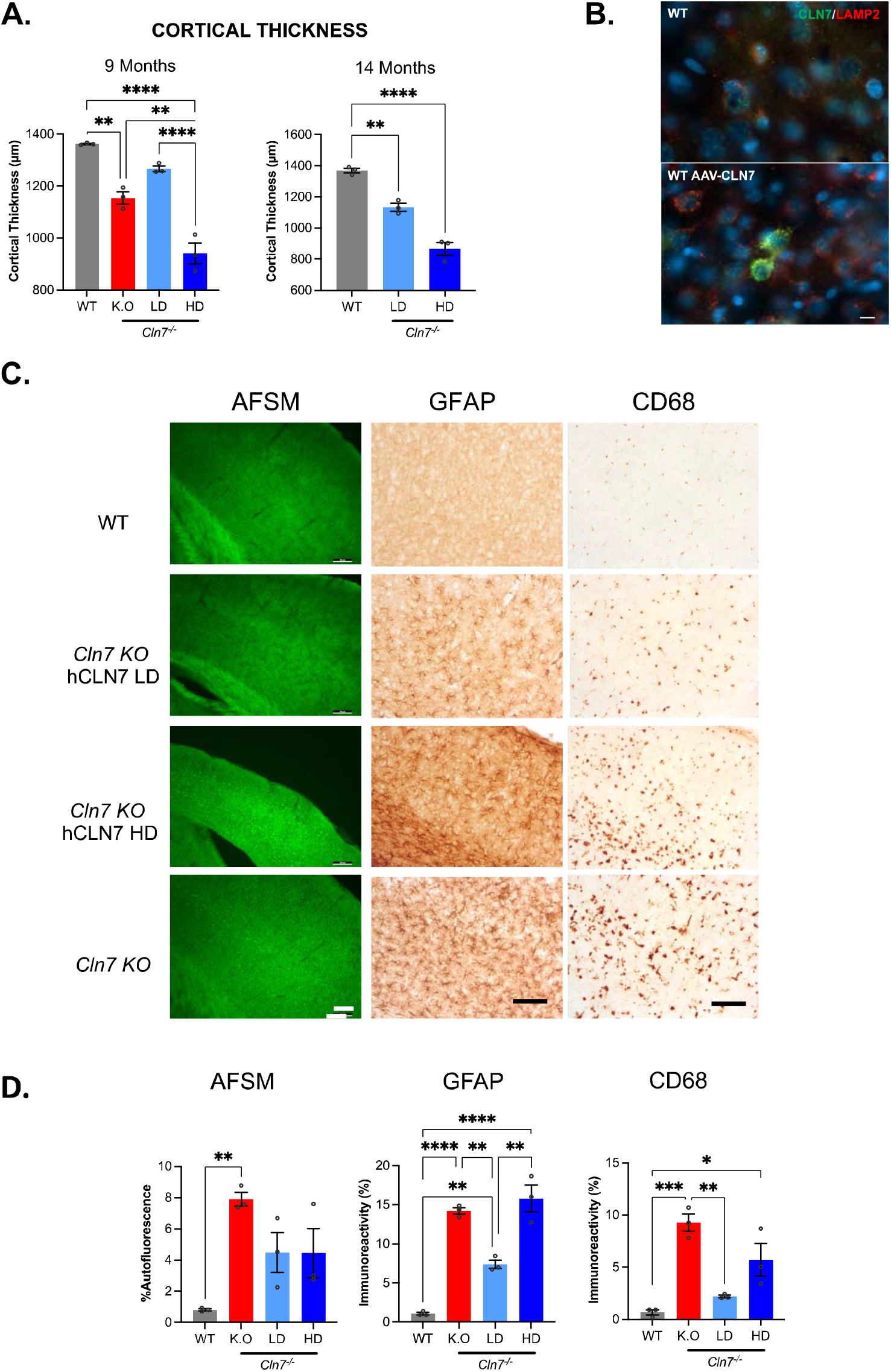
Dosage-dependent decreases in cortical thickness and elevated neuroinflammtion following intracerebroventricular AAV9-CLN7 gene therapy in *Cln7^Δex2^* mice. (**A**) Measures of cortical thickness of S1BF in NeuN-immunostained sections at 9 and 14 months, analyzed using a one-way ANOVA with post-hoc Bonferroni correction (n=3). (**B**) Representative images of CLN7 and LAMP2 immunohistochemistry of LD AAV-hCLN7 treated wild-type brain sections (x64 mag; Scale bar=10μm). (**C**) Representative images and (**D**) automated quantification of autofluorescent storage material (AFSM), Glial Fibrillary Acidic Protein (GFAP) as a marker of activated astrocytes, and CD68 as a marker of activated microglia in the S1BF cortex region at 9 months. Groups compared are wild-type (WT), untreated *Cln7^Δex2^* (KO), low-dose treated (LD) and high-dose treated (HD) *Cln7^Δex2^* mice. Scale bar=100μm, analyzed using a one-way ANOVA with post-hoc Bonferroni correction (n=3). All data presented as mean ± SEM, * p≤0.05, **≤0.01, ***≤0.001 and ****≤0.0001, (P-values in Supplementary Table 1).

### CLN7 iNPC present the same metabolic phenotype as the *Cln7^Δex2^* mouse

Much of the current understanding of CLN7 disease and subsequent therapeutic intervention is based on research conducted on the *Cln7^Δex2^* mouse model (*18*). We sought to explore a more clinically relevant model by generating iPSC from two patients with classic late infantile CLN7 disease. Pa380 is a female diagnosed at 2.5-years who was homozygous for the most frequent mutation c.881C>A causing a missense p.(Thr294Lys); this is a founder mutation within the Roma community (*21*). Pa474 is a male diagnosed at 4.5-years who was homozygous for c.1393C>T causing a missense p.(Arg465Trp) mutation (**Supp. Fig. 6A**). iPSC lines were generated from dermal fibroblasts archived at the BD repository held at UCL, London (https://www.ucl.ac.uk/ncl-disease/) and control iPSC generated from dermal fibroblasts from two 6-year-old donors (**Fig. 3A and Supp. Fig. 6B**). Following confirmation of genotype, iPSCs were subjected to targeted differentiation to nestin^+^ neural progenitor cells (iNPC), capable of maturation to βIII-tubulin expressing neurons, using a well-established protocol (*22*). After 30 days of neural maturation Pa474 neuronal cultures showed no overt disease phenotype. However, after 70d of extended culture we observed large aggregates accumulating in LAMP1^+^ vesicles indicative of enlarged lysosomes in Pa474 neurons (**Fig. 3B**). We established homogeneous proliferative cultures of nestin^+^ Pa380, Pa474 (collectively referred to as BD iNPC) and age-matched control iNPC maintained over multiple passages with little phenotypic variance (*22*) to evaluate as a cell model of CLN7 disease. Under steady-state culture conditions ICC showed that in control iNPC CLN7 was present in cytosolic vesicles and in the nucleus. This nuclear localization was unexpected but ICC using the same antibody, raised against the first 37 amino acids of human CLN7/MFSD8, has resulted in a similar cellular distribution in A431, U2OS and U251 cell lines analyzed by the Human Protein Atlas program (proteinatlas.org) and human dermal fibroblasts (*20*). In both BD iNPC, mutant CLN7 was expressed but was concentrated in larger perinuclear vesicles and relatively occluded from the nucleus (**Fig. 3C**).

**Figure 3.**
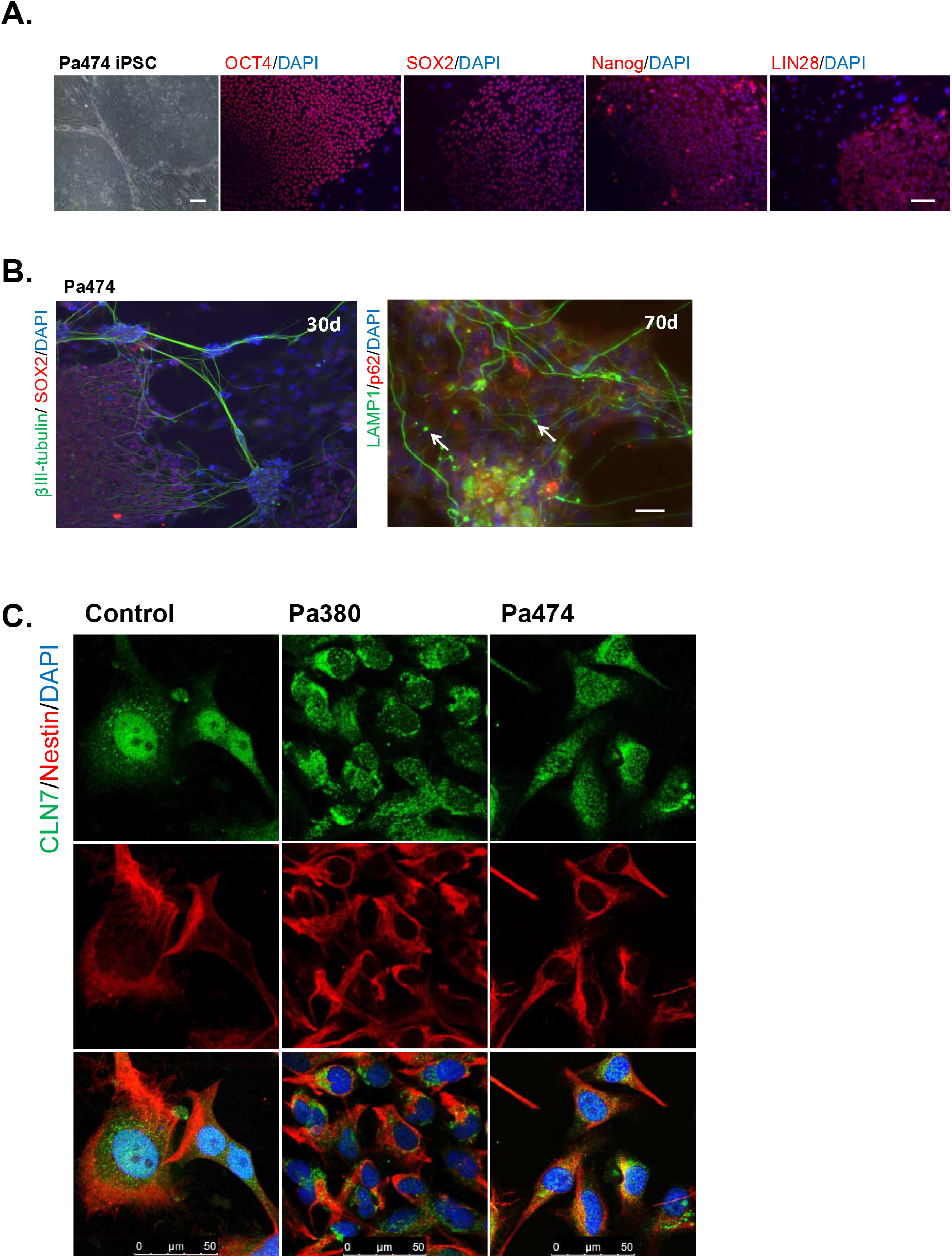
CLN7 patient-derived iNPC present a disease-specific cellular phenotype. (**A**) Induced pluripotent stem cell lines were generated from CLN7 patient Pa474 and immunocytochemistry (ICC) conducted for OCT4, SOX2, Nanog and LIN28 as indicators of transgene independent pluripotency (scale bars=100 μm). (**B**) ICC for markers of iNPC neural maturation; SOX2 and ßIII-tubulin and intracellular vesicles; p62 and LAMP1 during neural specification over 70 days (scale bar=50μm). White arrows indicate enlarged LAMP1^+^ vesicles. (**C**) ICC for CLN7 and the iNPC marker Nestin on control, Pa380 and Pa474 CLN7 patient iNPC (scale bar=50μm).

We recently described a metabolic phenotype in *Cln7^Δex2^* mice caused by mitochondrial dysfunction resulting in elevated ROS, bioenergetic collapse (*17*). In alignment with this study, BD iNPC had elevated mitochondrial membrane potential and superoxide production compared to control iNPC under steady-state conditions (**Fig. 4A & B**). In order to directly compare metabolic phenotype between BD iNPC and *Cln7^Δex2^* mice we isolated and cultured cortical neurons and conducted a MitoStress test. There was a nonsignificant trend for reduced mitochondrial ATP production in CLN7 iNPC as well as *Cln7^Δex2^* cortical neurons under steady-state conditions (**Fig. 4C, Supp. Fig. 7**). Bafilomycin A (Baf A) disrupts lysosomal acidification through inhibition of vATPase and also ER calcium reserves through SERCA inhibition (*23*). We hypothesized that Baf A treatment could exacerbate disease phenotype in BD iNPC by further compromising lysosomal function and elevating cytosolic calcium. Baf A treatment (100nM, 6h) resulted in a significant reduction in the ability of BD iNPC and *Cln7^Δex2^* mouse cortical neurons to respond to cellular stress by producing ATP through mitochondrial respiration when compared to controls (**Fig. 4C, Supp. Fig. 7**). Mitochondrial ATP production is quantified as the relative reduction in oxygen consumption rate after inhibition of the terminal complex V with oligomycin and hence disruption of the mitochondrial electron transport chain (ETC). Complex I of the ETC catalyzes the conversion of NADH to NAD+. NADH/NAD+ ratio was depleted in BD iNPC compared to control and, unlike control iNPC, was also unable to energetically respond to Baf A treatment (**Fig. 4D**). Furthermore, mRNA transcript for ATP5A, a mitochondrial protein which lies upstream of oligomycin inhibition, was elevated by ~10-fold in BD iNPC compared to control whereas UCP2 transcript, which is downstream of complex V, was decreased by <0.5-fold. A classical BD phenotype is the accumulation of lysosomal lipofuscin, rich in subunit c of mitochondrial ATP synthase (SCMAS) of which ATP5A is a component. It is entirely plausible that a gene regulatory feedback loop, the consequence of mitochondrial dysfunction, leads to mitophagy and the aberrant accumulation of SCMAS at the dysfunctional lysosome. Our data are consistent with Baf A exacerbating a BD phenotype in *Cln7^Δex2^* mouse cortical neurons and CLN7 patient iNPCs that links lysosomal and mitochondrial defects.

**Figure 4.**
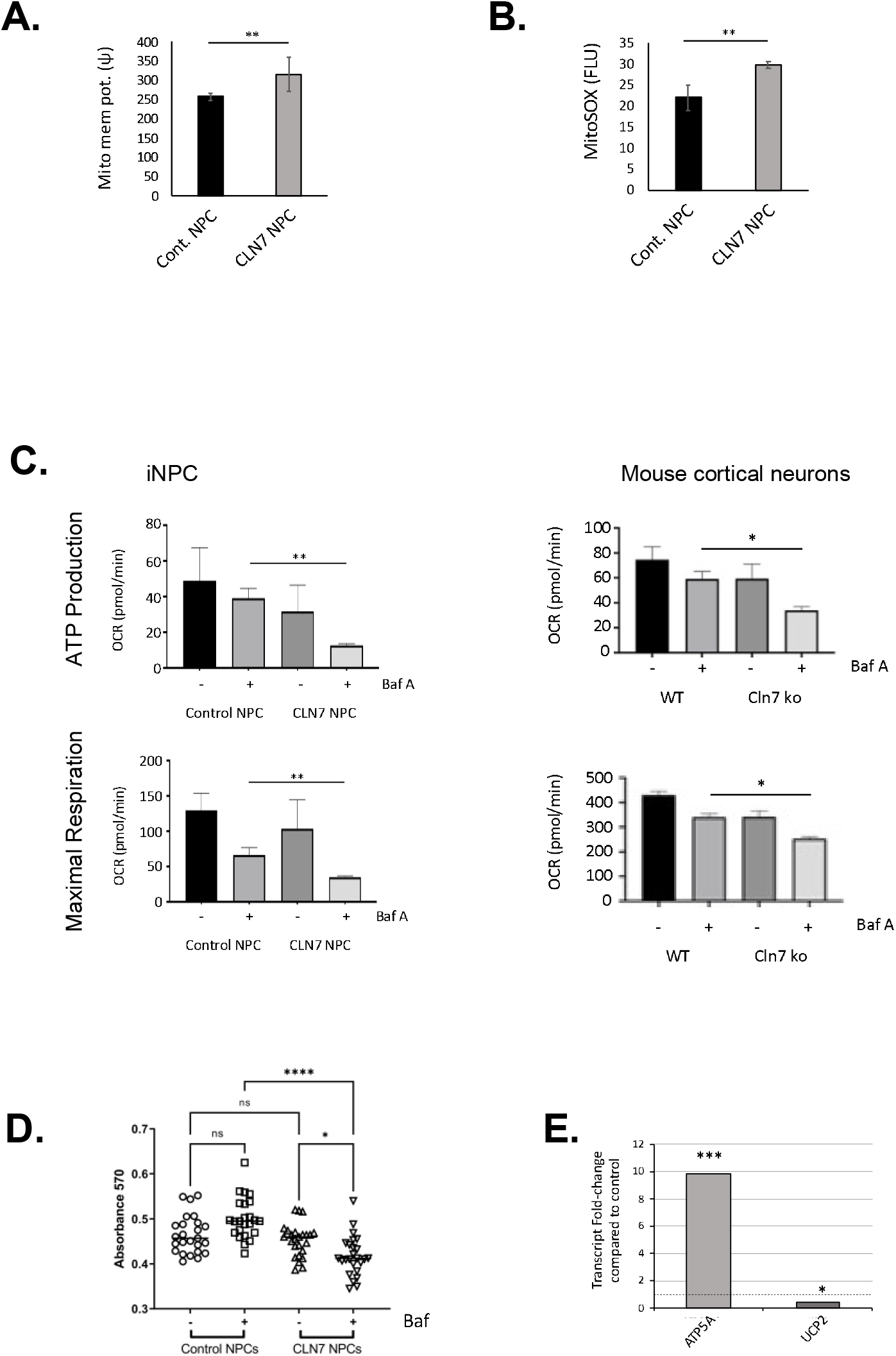
CLN7 iNPC and primary *Cln7^Δex2^* cortical neurons share a common metabolic disease phenotype that is exacerbated by Baf A. (**A**) Mitochondrial membrane potential measured by TMRM and (**B**) mitochondrial superoxide measured using MiotoSOX assay in CLN7 compared to control iNPC (n=6). (**C**) Mitochondrial respiration assayed in the presence or absence of Baf A in CLN7 versus control iNPC and *Cln7^Δex2^* versus wild-type cortical neurons. Change in oxygen consumption rate (OCR) quantified before and after addition of oligomycin for ATP production then FCCP followed by rotenone/antimycin A for maximal respiration in the presence or absence of Baf A in CLN7 versus control iNPC (n=6) and in *Cln7^Δex2^* versus wild-type cortical neurons (n=4). (**D**) Quantitation of NADH/NAD+ ratio by MTT assay in the presence and absence of Baf A in CLN7 compared to control iNPC (n=6). Quantitation of (**E**) ATP5A and UCP2 transcripts by qRT-PCR (n=4). Baf A treatment was 10nM for 6h, error bars indicate S.D., pairwise comparisons were made using a one-tailed t-test. All data presented as mean ± SEM, * p≤0.05, **≤0.01, ***≤0.001 and ****≤0.0001, (P-values in Supplementary Table 1).

### CLN7 sub-cellular redistribution in normal and BD iNPC following Baf A treatment

Having identified differential vesicular and nuclear localization of CLN7 between control and BD iNPC (**Fig. 3C**) we compared its cellular distribution under steady-state culture conditions and after incubation with Baf A. BafA inhibits the acidification of the lysosome and has been reported to block the fusion of autophagosomes with lysosomes. Under basal conditions CLN7 co-localized with some but not all p62^+^ large perinuclear vesicles. Interestingly, there were far fewer and smaller p62^+^ vesicles in both BD iNPC with mutant CLN7 concentrated in small perinuclear vesicles (**Fig. 5A**). Following 6h Baf A treatment (100nM, 6h) there was a rapid redistribution of CLN7 protein in iNPC. CLN7 spreads throughout the cell following Baf A treatment in control iNPC, including the nucleus, but cytosolic distribution did not appear vesicular or co-stain with p62. In complete contrast mutant CLN7 translocates to the nucleus following Baf A treatment in BD iNPC (**Fig. 5B**). This was consistent between Pa380 and Pa474 iNPC with their different disease-causing missense mutations in exons 10 and 13 respectively (**Supp. Fig. 6**). Additionally, we noticed massively elevated p62 immunostaining and higher levels of cell death in CLN7 iNPC. Under steady-state conditions, apoptosis, as quantified by Apotracker staining on viable cells, was significantly higher in Pa380 and Pa474 iNPC compared to control. There was no increase in apoptosis after Baf A treatment in control iNPC but it further elevated apoptosis in Pa380 and Pa474 BD iNPC (**Fig. 5C**). Here, for the first time, we prove that disease-causing missense mutations in CLN7 do not prevent expression of a protein that is conditionally responsive, albeit in an aberrant manner consistent across two genotypes. One possible inference is that genetic mutation results in a malevolent protein gain-of-function phenotype as well as a probable canonical loss-of-function, this could ultimately lead to neuronal apoptosis through an undefined mechanism.

**Figure 5.**
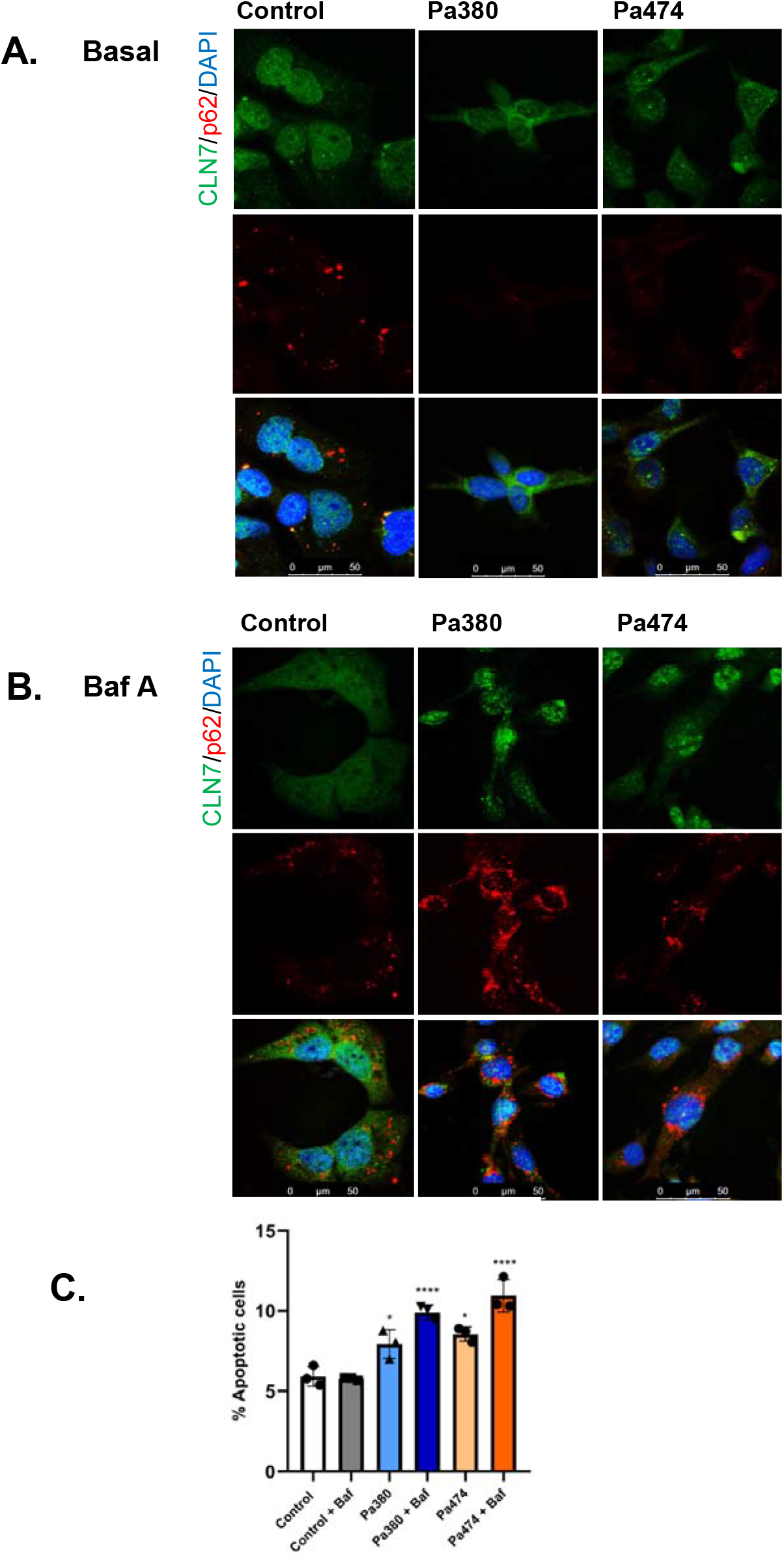
Baf A treatment of iNPCs causes a rapid cellular redistribution of CLN7 protein that is altered by disease. Intracellular localization by ICC for CLN7 and p62 in normal and Pa380 and P474 CLN7 patient iNPC with DAPI nuclear counterstain in the (**A**) absence and (**B**) presence of Baf A (representative images, n=3. Size bars=50μm). (**C**) Apoptosis assessment using Apotracker staining of gated viable cells that exclude eFluor-780 dye in the absence and presence of Baf A (10nM for 6h), error bars indicate S.D., pairwise comparisons were made using a one-tailed t-test. All data presented as mean ± SEM, * p0.05, **≤0.01, ***≤0.001 and ****≤0.0001, (P-values in Supplementary Table 1).

### Proteomics reveal a profound nuclear defect in CLN7 iNPC

A global quantitative proteomic profiling was conducted comparing Pa380 and Pa474 with control iNPCs from two age-matched healthy donors after Baf A treatment or vehicle control. Experiments were performed in triplicate biological repeats for each individual and condition. Three separate mass spectrometry runs used three separate 8-plexes of isobaric tags for absolute and relative quantification (iTRAQ). A total of 24 iTRAQ tags were used to label the prepared peptides of the 24 samples so that each proteomic quantification run had the peptide sample of each cell sample with and without Baf A treatment (**Fig. 6A, Supp. Fig. 8**). A comprehensive Bayesian analysis with pairwise comparisons was conducted as shown in **Table 1** with a total of 5619 proteins quantified and compared across all samples. Differentially regulated proteins were analyzed for pathway changes using Reactome Pathway application through Cytoscape and for cellular components using gProfiler through Cytoscape (*24*). The top Reactome terms listed alongside the number of proteins within class and a combined sliding scale of p value statistical significance with a cut-off of p<0.05. Under basal conditions proteins associated with membrane transport were strongly upregulated in BD iNPC compared to control; these largely localize to the cytosolic and endomembrane compartments (**Fig. 6Bi, Supp. Fig. 9A**). When pathways were expanded to look at minor divisions Intra-Golgi and ER-Golgi COPI/II transport was the highest enriched subdivision. The second highest enriched major pathway was Asparagine N-linked glycosylation integral to post-translational modification for proteins synthesized and folded in the ER. Most upregulated proteins could be affiliated with a cellular stress response, and interestingly 13 proteins of the PSMD family, subunits of 26s proteasome, were all significantly upregulated suggesting highly active proteostasis. In stark contrast, the vast majority of proteins downregulated in BD compared to control iNPC under basal conditions were nuclear in localization and functions include rRNA processing, mRNA processing and splicing and transcription. Of 599 significantly downregulated proteins, 442 (74%) were functionally nuclear (**Fig. 6Bii, Supp. Fig. 9B**). These data imply a pan-nuclear defect that would be consistent with a nuclear import/export defect in CLN7 BD. A targeted analysis showed that 23 nuclear pore complex associated proteins were downregulated in our dataset (**Fig 6Biii**).

**Figure 6.**
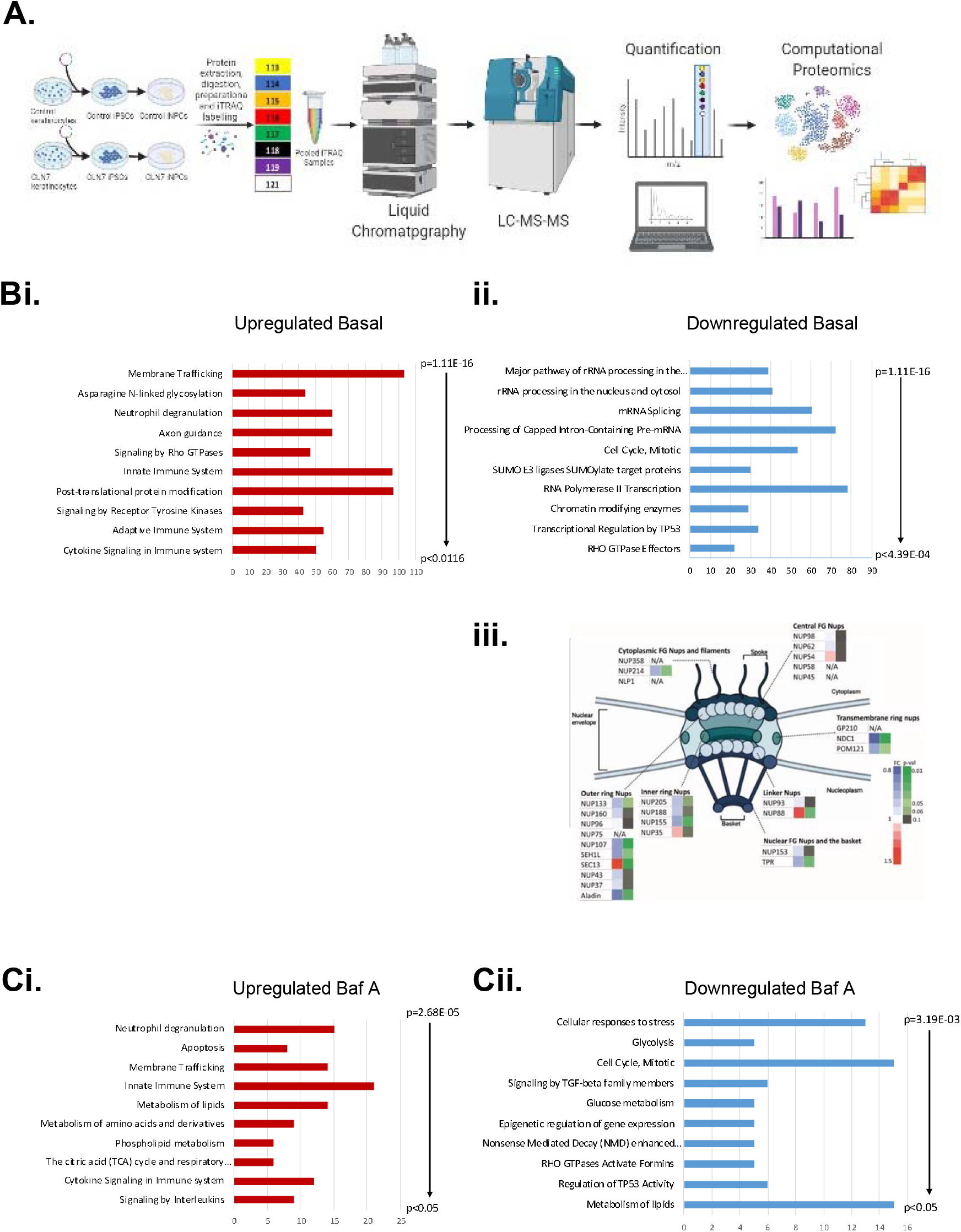
Unbiased global proteomics analysis reveals a unique nuclear defect in CLN7 iNPC. (**A**) Schematic workflow of iTRAQ mass spectrometry, proteomic quantification and bioinformatic analyses. A total of 5619 proteins were compared for alterations in expression between combined Pa380 and Pa474 and two age-matched control iNPC under steady-state conditions and following Baf A treatment. Samples were analyzed by hierarchical Bayesian modelling then pathway changes identified using Cytoscape gProfiler through Reactome Pathway. (**B**) Reactome terms are listed on y-axes ranked by p-value with a cut-off of p<0.05 and the number of deregulated proteins within that term on the x-axes. Terms. Upregulated (**i**) and downregulated (**ii**) proteins in CLN7 compared to control iNPC with (**iii**) schematic representation of the nuclear pore complex identifying deregulated proteins. (**C**) Upregulated (**i**) and downregulated (**ii**) proteins in CLN7 compared to control iNPC after Baf A treatment. Statistical evaluations are detailed in the Methods section.

**Table 1.** Numbers of proteins deregulated in group pairwise comparisons

| Comparison | Proteins with sig. q-value | Proteins upregulated (20% cut-off) | Proteins downregulated (20% cut-off) |
| --- | --- | --- | --- |
| Control iNPC vs Control iNPC (Baf A+) | 0 | - | - |
| BD iNPC vs BD iNPCs (Baf A+) | 45 | 0 | 1 |
| Control iNPC vs BD iNPC | 1366 | 762 | 599 |
| Control iNPC vs BD iNPC (Baf A+) | 386 | 147 | 238 |

In equivalent comparisons following Baf A treatment, similar membrane trafficking pathways were upregulated in BD compared to control iNPC confirming experimental continuity. However, the second most significant Reactome term was now “Apoptosis”. In addition, metabolic pathways such as the citric acid (TCA) cycle and respiratory electron transport, phospholipid metabolism, metabolism of amino acids and derivatives and metabolism of lipids, were among the most significant pathway changes with the highest number of upregulated proteins (**Fig. 6Ci, Supp. Fig. 9C**). As well as a substantial proportion of nuclear proteins downregulated (166 of 238; 70%) after Baf A treatment, glycolysis and metabolism of glucose and lipids are also strongly represented (**Fig. 6Cii, Supp. Fig. 9D**).

To identify key proteins that lie at a nexus where protein-protein signaling can be substantially impaired, minimal essential networks (MEN) were built for our four comparison networks based on the public UniProt data base through Cytoscape (**Supp. Fig. 10**). MEN contain the top 10% of interactome proteins scored for connectivity and nexus network properties which represent the most functionally related regions of interactome models (*25*). From the UniProt networks the top 10% of “hubs” and “bottlenecks” within the network were identified using the Cytohubba algorithm and used to generate MEN as shown in **Table 2** where dominator nodes are listed for each network. VDAC1 is the exclusive dominator to upregulated networks in BD iNPCs compared to controls. While PGAM5, UCHL5 and HSPA8 are dominators to downregulated networks in CLN7 BD iNPCs. Intriguingly HSPA8 (also known as HSC70) and PGAM5 both mechanistically stabilize PINK1, whilst VDAC1 is a target for Parkin-mediated ubiquitylation during ROS-induced mitophagy (*26*–*28*). UCLH5 is a deubiquitinase that elevates proteasomal proteostasis (*29*). Collectively, these data correlate well with an emerging hypothesis of lysosomal and nuclear defects combining to destabilize cellular metabolism, energy deficit and ultimately resulting in elevated mitophagy and mitochondrial-mediated apoptosis.

**Table 2.** Minimal essential network node proteins from CLN7 v control comparisons

| Minimal Essential Network (MEN) Nodes |  |  |  |  |  |
| --- | --- | --- | --- | --- | --- |
| Upregulated Basal | PRNP | ZDHHC5 | Vps29 | Rala | RALY |
|  | GCN1 | **VDAC1** | TLN2 | FUS | MAPT |
| Downregulated Basal | FAF1 | PRNP | ACTL6A | THRAP3 | SMARCE1 |
|  | TP53BP1 | DDX42 | ZDHHC5 | METTL3 | MAPT |
| Upregulated Baf A | PRNP | FUS | ZDHHC5 | SLC16A11 | MAPT |
|  | Vps29 | **VDAC1** | GCN1 | RALY | TJP1 |
| Downregulated Baf A | SLC16A11 | FAF1 | HSP90AA1 | METTL14 | **PGAM5** |
|  | HMGB1 | **HSPA8** | PGAM5 | **UCHL5** |  |

### CLN7 exists as multiple isoforms

CLN7 is ubiquitously expressed (gtexportal.org) but genetic mutation results overwhelmingly in neurodegenerative disease. Analysis of publicly accessible RNA-seq data collected on 54 tissues from nearly 1000 individuals through the GTEx Portal (gtexportal.org) shows that CLN7 (MFSD8: ENSG00000164073.9) is expressed in all tissues but is highest in brain cerebellum (**Supp. Fig. 11**). CLN7 expression was evaluated by western blot in HEK293T, SH-SY5Y and MEF lysates and a variable array of up to 11 reproducible bands ranging from ~22kDa to ~180kDa were detected (**Fig. 7A**). The full-length CLN7 cDNA codes for a 518 amino acid protein of approximately 57kDa that has been experimentally validated in other cells (*30*). A similar profile of western blot bands was previously identified using an antibody raised against amino acids 246-266 of CLN7 in wild-type MEFs (*31*). We assume that the larger bands are the result of dimerization and/or protein modification but we did not detect any differences between denaturing/reducing and non-denaturing/non-reducing conditions for western blot (data not shown). The structure of CLN7 remains experimentally unresolved but other MFS family proteins such as the GLUTs are known to form functionally distinct homotetramers (*32*) so this is a plausible rationale for larger bands on western blot but remains to be explored. However, we did note in SH-SY5Y neuroblastoma cells that a ~60kDa band was susceptible to peptide N-glycosidase F (PGNase F) digestion with a concomitant dose increase in the ~57kDa band. We conclude that full-length CLN7 is the full-length N-glycosylated form (~60kDa) resulting in a size shift on PAGE (**Fig. 7Bi**). It is intriguing that the N-glycosylated form is not evident in iNPC; Steenhuis *et al*. previously described two N-glycosylation sites at amino acids 371 and 376, incorporated into the luminal loop L11, of full-length CLN7 (*30*). Neither the presence or relative abundance of isoforms was affected by incubation with Baf A or the more specific SERCA inhibitor thapsigargin in SH-SY5Y cells implying that the changes in sub-cellular localization do not reflect either gene expression or protein modification steps (**Fig. 7Bii**).

**Figure 7.**
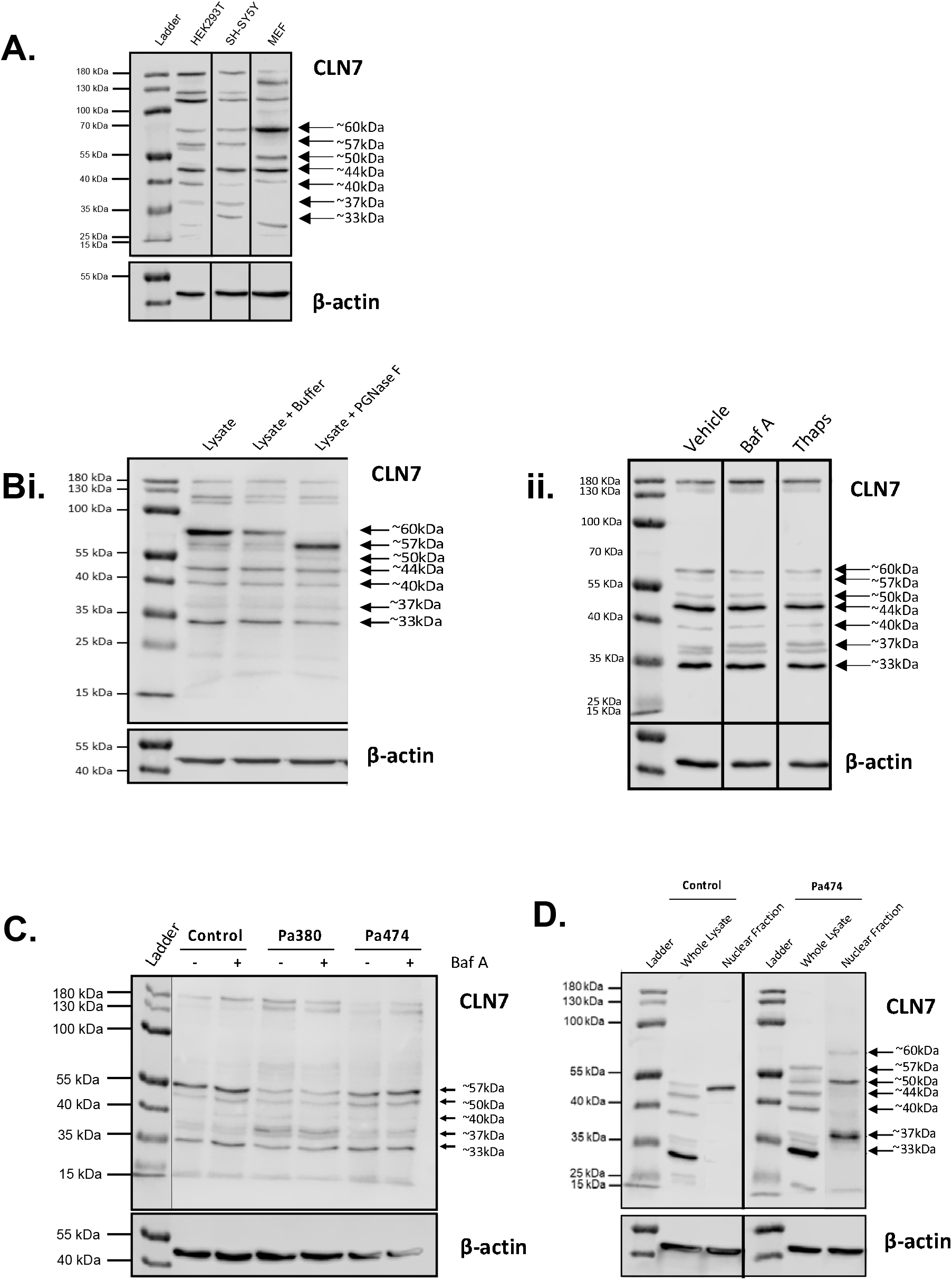
CLN7 has multiple isoforms with differential localization in CLN7 iNPC. CLN7 western blots conducted on lysates from; (**A**) HEK293T, SH-SY5Y and wild-type MEFs; SH-SY5Y in the presence or absence of (**Bi**) PNGase F, (**Bii**) Baf A or Thapsigargin. (**C**) CLN7 western blots conducted on lysates from CLN7 (Pa380 and Pa474) and control iNPC in the presence and absence of Baf A, representative of 3 biological repeats (Supp Fig. 12). (**D**) CLN7 western blots conducted on whole cell lysates and nuclear fractions from CLN7 (Pa474) and control iNPC in the presence and absence of Baf A, representative of 3 biological repeats (Supp Fig. 12).

Protein lysates from BD and control iNPC were evaluated over multiple biological repeats and western blots for changes in CLN7 expression profile (**Fig. 7C and Supp Fig. 12A**). Overall, we did not detect any reproducible alterations in band intensities between BD and control iNPC with or without Baf A treatment. This would suggest that sub-cellular redistribution of CLN7 in response to Baf A treatment is not driven at the level of gene expression, splicing or translation initiation but could be due to protein modification or conditional redistribution of existing CLN7 isoforms. Others have established that the 57kDa isoform of CLN7 localizes to the lysosomal membrane (*18*, *33*). Based on our confirmation of CLN7 nuclear localization using ICC we fractionated nuclei from BD and control iNPCs and compared with CLN7 expression of whole lysates on western blot. A ~50kDa band was evident in nuclear fractions from control and BD iNPC but a ~37kDa band was present only in BD iNPC (**Fig. 7D**) across multiple biological repeats (**Supp. Fig. 12B**). In combination with alterations in cellular distribution, the existence of this ~37kDa CLN7 nuclear isoform are novel unique identifiers of CLN7 BD in patient iNPC.

### CLN7 transgene isoforms localize to the nucleus

Having established that CLN7 is expressed in multiple isoforms we sought to rationalize the band sizes from western blot with predicted isoforms generated through post-transcriptional and post-translational modifications. There are 62 potential CLN7 transcript variants annotated on Ensembl across the 13 exons with 24 predicted as protein coding. Of interest is transcript ENST00000641690.1, generated by splice excision of exons 7 and 8. This results in a putative 451 amino acid 50 kDa protein (CLN7^Δex7/8^) consistent with the size of the isoform detected in nuclear fractions. Steenhuis *et al*. previously reported that CLN7 is susceptible to proteolytic cleavage at luminal loop L11 which contains two cysteine residues Cys^401^ and Cys^408^. Assuming cleavage occurs due to lysosomal cathepsin cleavage, then cathepsins L and C are predicted to cleave CLN7 between the C^L residues (https://www.ebi.ac.uk/merops/index.shtml; **Supp Fig. 12C**). Protease cleavage of CLN7 at Cys^401^ results in a putative 400 amino acid 44kDa isoform (CLN7^Cys401^) but intriguingly protease cleavage at the same Cys residue in CLN7^Δex7/8^ results in a 37kDa isoform (CLN7^Δex7/8-Cys401^) consistent with that found in the nuclear fraction of BD iNPC (**Table 3**). We generated expression plasmids containing the human coding sequence for the full-length isoform and each of these three truncated isoforms (**Fig. 8A**) to evaluate cellular localization in transfected cells. First, we transfected wild-type and *Cln7^Δex2^* mouse cortical neurons with an expression plasmid encoding the full-length GFP-CLN7 fusion transgene (**Fig. 8B**). As with endogenous CLN7 protein, GFP-CLN7 partially co-localized with LAMP1 and p62 in WT neurons. In *Cln7^Δex2^* neurons there was considerable co-localization of GFP-CLN7 and p62 in enlarged perinuclear vesicles but very little GFP-CLN7/LAMP1 co-localization, challenging the notion that CLN7 localizes only to lysosomal membranes. Most notably, there was no evidence of nuclear localization of GFP-CLN7 in either wild-type or *Cln7^Δex2^* neurons strongly suggesting that full-length GFP-CLN7 neither localizes to the nucleus nor is capable of being post-translationally modified to a nuclear localizing isoform.

**Figure 8.**
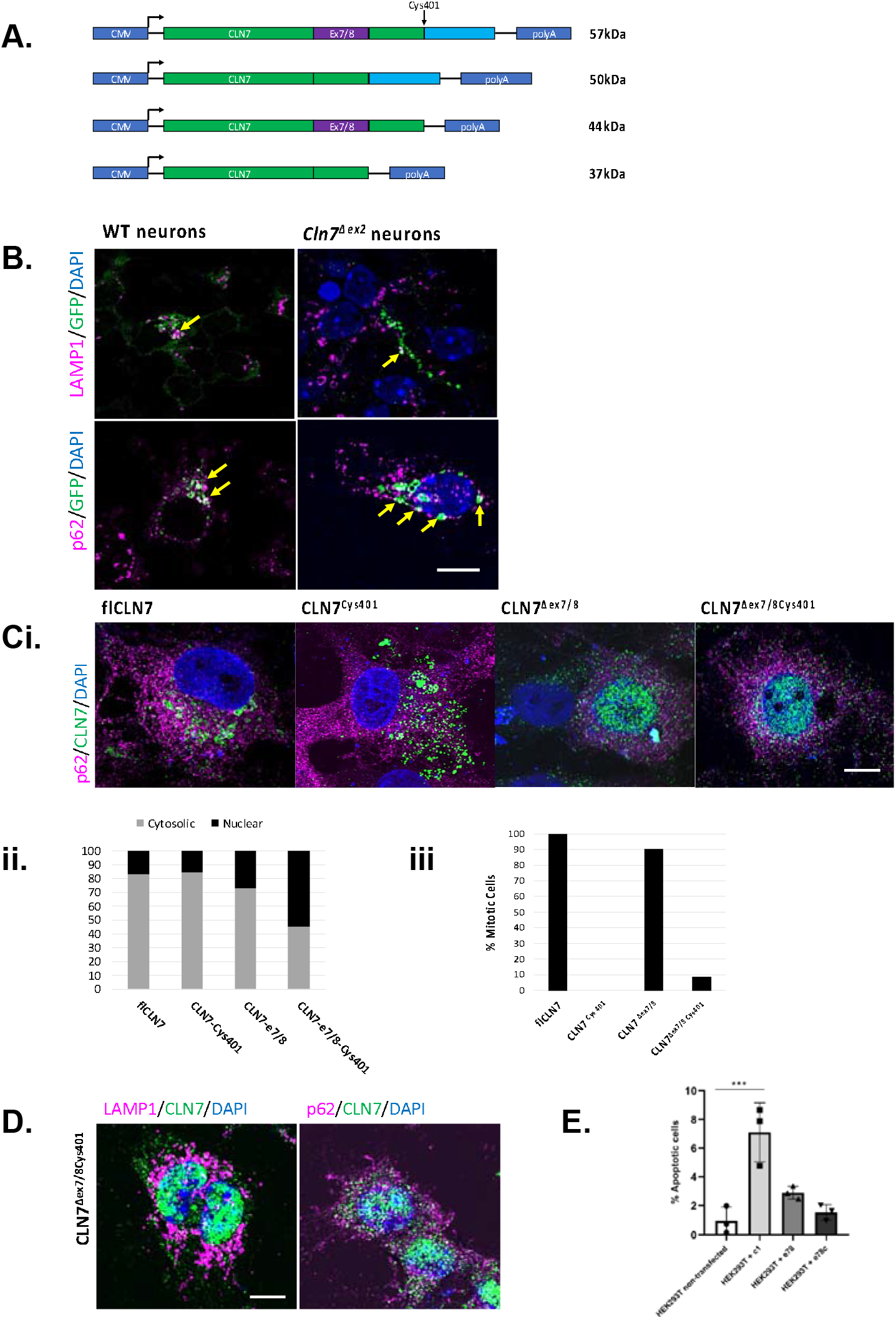
Transgenic CLN7 isoforms differentially localize in *Cln7^Δex2^* cells and increase apoptosis. (**A**) Schematic representation of full-length and truncated gene variants of CLN7. (**B**) ICC for CLN7, LAMP1 and p62 with DAPI nuclear counterstain following transfection of WT and *Cln7^Δex2^* cortical neurons with a GFP-CLN7 fusion construct. Yellow arrows indicate areas of co-staining. (**Ci**) ICC for CLN7 and p62 with DAPI nuclear counterstain following transfection of *Cln7^Δex2^* MEFs with CLN7 isoform constructs with (**Cii**) quantitation of nuclear and cytosolic localization (n=30 cells) (**Ciii**) Quantitation of CLN7 positive transfected cells undergoing mitosis (n=30 cells). (**D**) ICC for CLN7, LAMP1 and p62 with DAPI nuclear counterstain following transfection of *Cln7^Δex2^* cortical neurons with the CLN7^Δex7/8-Cys401^ expression plasmid. (E) Apotracker staining of gated viable cells that exclude eFluor-780 dye following transfection of HEK293T cells with the CLN7 expression plasmids. Error bars indicate S.D., pairwise comparisons were made using a one-tailed t-test. All data presented as mean ± SEM, * p0.05, **≤0.01, ***≤0.001 and ****≤0.0001, (P-values in Supplementary Table 1). All images are representative of 3 independent experiments, size bars=50μm.

**Table 3.** Putative CLN7 isoforms predicted from mRNA splicing and cysteine protease digestion

| Size (kDa) | Putative CLN7 isoform | Cont. Lysate | Cont. Nuclei | CLN7 Lysate | CLN7 Nuclei |
| --- | --- | --- | --- | --- | --- |
| 60 | Full-length N-glycosylated |  |  | ✓ | ✓ |
| 57 | Full-length | ✓ |  | ✓ |  |
| 50 | Transcript variant Δex7/8 | ✓ | ✓ | ✓ | ✓ |
| 44 | Cys401 protease digested | ✓ |  | ✓ |  |
| 40 | Transcript variant Δex12/13 | ✓ | | ✓ | |
| 37 | Transcript variant Δex7/8 Cys401 | | | | ✓ |
| 33 | Transcript variant Δex6/7/8 Cys401 | ✓ | | ✓ | |

We transfected *Cln7^Δex2^* mouse embryonic fibroblasts (MEF) with expression plasmids encoding full-length (fl), or truncated isoforms of CLN7 and assessed sub-cellular localization in reference to p62 vesicles. Similar to endogenous and GFP-CLN7, flCLN7 showed partial overlap with p62 vesicles, as did CLN7^Cys401^, both of which presented as largely vesicular and being largely but not exclusively excluded from the nucleus. However, CLN7^Δex7/8^ and CLN7^Aex7/8-Cys401^ expression was distributed between the cytosolic vesicles and the nucleus (**Fig. 8C**). Quantitation of nuclear versus cytosolic immunoreactivity showed 54% of CLN7 was nuclear in CLN7^Aex7/8-Cys401^ compared to 27% with CLN7^Δex7/8^ and 15-20% in CLN7^Cys401^ and flCLN7 (**Fig. 8Cii**). Interestingly, 90-100% of the flCLN7 and CLN7^Δex7/8^ transfected MEFs that had CLN7 immunoreactivity in the nucleus were actively undergoing mitosis, something that would not occur in post-mitotic neurons (**Fig. 8Ciii**). This may imply there is an unrecognized role of CLN7 in cell division, but this requires further investigation. Having identified the 37kDa CLN7^Aex7/8-Cys401^ isoform in CLN7 patient iNPC nuclei we transfected *Cln7^Δex2^* mouse cortical neurons with this expression vector and confirmed nuclear localization and an absence of immunolocalization with LAMP1 or p62 vesicles (**Fig. 8D**). Our data imply that multiple CLN7 isoforms have the capacity to enter the nucleus in proliferative cells but only CLN7^Aex7/8-Cys401^ accumulates in the nucleus in non-mitotic cells. Finally, we assessed apoptosis in transfected viable HEK293T cells and found that all three isoforms induced significantly higher levels of apoptosis than a plasmid expressing GFP but interestingly, the CLN7^Cys401^ isoform caused the highest level of apoptosis (**Fig. 8F**). Overall, our data imply that CLN7 isoforms are spatially and functionally distinct, influenced both by cellular stress and cell cycle. Together, the data presented here suggest that disease-causing mutations in CLN7 may result in loss-of-function but also a disruptive gain-of-function that contributes to disease.

## Discussion

Like many monogenic neurodegenerative diseases, gene complementation using high-titer AAV vectors is a compelling therapeutic option for most types of Batten disease including CLN7. However, AAV vectors have limited payload capacity, meaning the expression cassette is most often limited to a strong minimal promoter driving expression of a cDNA with the exclusion of gene regulatory elements. Such an approach makes the dual assumption that; (a) disease is caused exclusively by loss of protein function encoded by the full-length, unmodified cDNA and, (b) elevated levels of transgenic protein are not toxic. We conducted neonatal intracerebroventricular administration of an AAV-hCLN7 gene therapy in *Cln7^Δex2^* knockout mice, a well characterized and well used model of CLN7 disease (*18*). Gene therapy resulted in extended lifespan in *Cln7^Δex2^* mice. The expressed CLN7 transgenic protein is localized to LAMP2 vesicles and there is a reduction in lysosomal AFSM, known to pathologically accumulate in BD neurons. *Cln7^Δex2^* mice present significant cortical shrinkage at 9-months compared to wild-type. Cortical shrinkage was partially rescued with LD but significantly exacerbated by HD AAV-hCLN7 gene therapy. This was mirrored by poorer performance in a battery of locomotor function and behavioral tests in HD compared to LD AAV-hCLN7 gene therapy. Concomitantly, markers of astrocyte (GFAP) and microglial (CD68) activation were reduced in LD but unchanged in HD compared to untreated *Cln7^Δex2^* mice. These data were mirrored at 14-months where reliable comparisons with untreated *Cln7^Δex2^* mice was not possible due to mortality. Collectively, our pre-clinical evaluation of AAV-hCLN7 in the well-established *Cln7^Δex2^* mouse model show indications of efficacy, particularly lifespan extension, but with clear evidence of dose escalating toxicity. Our data are in broad alignment with the recently published pre-clinical dataset of Chen *et al*. who administered similar doses of a similar vector to *Cln7^Δex2^* mice, albeit by intrathecal administration at P7-10 (*10*). They reported significant increases in survival and rotarod latency to fall at 9-months in LD and HD groups with therapeutic trends in other behavioral and neuroinflammatory evaluations that did not reach significance. However, our data shows statistically rigorous evidence of neurotoxicity and neuroinflammation relating to escalating dose of AAV-hCLN7 and expression driven by the powerful synapsin promoter. This is relevant for clinical trial design.

The unexpected gene therapy results provided the paradigm for a more in-depth study of CLN7 fundamental biology in relation to normal function. The *Cln7^Δex2^* knockout mouse is well characterized to present complete loss of CLN7 function. However, data from the UCL NCL repository (https://www.ucl.ac.uk/ncl-disease/) (*13*) indicates that for CLN7 at least 35/51 (69%) of disease-causing mutations are either missense or predicted splicing defects, with missense mutations representing nearly half of the known mutations and by far the most frequent founder mutations within regional communities worldwide. The single most frequent CLN7 disease allele is c.881C>A p.(Thr294Lys) so we generated iPSC from dermal fibroblasts from a patient (Pa380) homozygous for this mutation alongside another patient, Pa474 harboring a different missense mutation, c.1393C>T p.(Arg465Trp) in homozygosity. Patients with these mutations in homozygosity always have late infantile CLN7 disease. Neither mutation prevents correct transport to the lysosome in COS-7 cells (*30*). CLN7 and age-matched control iPSc were specified to iNPC capable of generating post-mitotic neurons. CLN7 neurons presented with enlarged lysosomes, and we have previously reported elevated SCMAS and disease-specific phenotypes in these CLN7 iNPC (*16*, *17*). Using a commercially available polyclonal antibody raised against the N-terminal 37 amino acids of CLN7/MFSD8, we detected characteristic immunostaining of cytosolic vesicles with partial overlap with LAMP1 and p62 immunoreactivity and intense staining of the nucleus. This is consistent with protein atlas data (https://www.proteinatlas.org/ENSG00000164073-MFSD8/subcellular#human) and observations previously reported by Geier *et al*. in human dermal fibroblasts (*20*). Importantly, both Pa380 and Pa474 iNPC showed immunoreactivity for CLN7 but the sub-cellular localization contrasted with that in controls, being largely perinuclear and occluded from the nucleus.

We have previously reported that CLN7 iNPC and *Cln7^Δex2^* mouse cortical neurons share a mitochondrial bioenergetic phenotype (*17*) so we sought to delineate what disease phenotypes in iNPC are due to loss-of-function in order to identify potential novel gain-of-function phenotypes. First, we confirmed that CLN7 iNPC have elevated mitochondrial membrane potential and ROS, consistent with mitochondrial stress. We evaluated mitochondrial bioenergetics following Baf A treatment, an inhibitor of vATPase and the ER Ca^2+^ channel SERCA (*34*). We found that discrete deficits in energy production between BD and control were massively exacerbated by Baf A *Cln7^Δex2^* mouse cortical neurons and phenocopied in CLN7 iNPC implying a common loss-of-function mechanism. The recent publication by Wang *et al*. describes CLN7 as being a chloride channel that sits at the membrane of autophagic-lysosomal vesicles, is activated by lysosomal acidification, and may regulate lysosomal Ca^2+^ efflux to the cytosol (*7*). This is consistent with exacerbation of the deregulation of Ca^2+^ distribution by CLN7 mutation and SERCA inhibition synergistically negatively affecting mitochondrial bioenergetics, possibly even through mitochondrial-ER contact sites (*35*). Further, Wang *et al*. present data showing that p.(Thr294Lys), the mutation carried by Pa380, causes defects in CLN7 chloride channel activity and the maturation of p62^+^ endolysosomes. Again, this is consistent with our own observations where, in control cells, we see co-localization of CLN7 more so with p62^+^ than LAMP^+^ vesicles. Baf A treatment for 6h results in the redistribution of CLN7 in control cells where it appears to become cytosolic but, unexpectedly, in CLN7 iNPC there is a pronounced nuclear localization and complete lack of p62 co-staining. Such a rapid redistribution is highly unlikely to be due to alterations in gene expression and more likely due to a dynamic cellular response to drug. We further noted that CLN7 iNPC were more susceptible to apoptosis under steady state culture and that this is exacerbated in both Pa380 and Pa474 iNPC following Baf A treatment. This implies a link between a stress-induced CLN7 defect linking vesicular transport, mitochondrial bioenergetics, nuclear function and induction of apoptosis.

We applied global, non-biased iTRAQ proteomics to identify deregulated pathways in CLN7 iNPC compared to controls under steady-state culture conditions and following Baf A treatment. A basal CLN7 disease phenotype includes elevated vesicular transport, particularly COPI/II ER-Golgi transport, and components of the 26s proteasome, all indicative of a cellular stress response and drive to maintain proteostasis. The most profound change noted is that 74% of significantly downregulated proteins are functionally resident in the nucleus, encompassing all major nuclear activities. There is a pronounced enrichment of downregulated proteins around the nuclear pore complex where obstruction of nuclear transport would be consistent with an overall depletion of nuclear proteins. Following Baf A treatment additional upregulated pathways include respiratory electron transport and TCA cycle consistent with a crisis in mitochondrial respiration concomitant with downregulation of glucose metabolism. Notably, the term “Apoptosis” is the second most significant upregulated pathway following Baf A treatment including key pro-apoptotic mediators DIABLO, cytochrome C and caspase-3, all involved in mitochondrial induced apoptosis. When we evaluated minimal essential networks (MEN) within our datasets we identified VDAC1 as a dominator of upregulated networks and PGAM5 and HSPA8 as dominators of downregulated networks in CLN7 disease. Intriguingly, all three are involved in quality control of mitochondria through Parkin/PINK1 mitophagy (*26*–*28*). We have discovered a novel nuclear defect in CLN7 disease as well as further defining vesicular and mitochondrial defects. Although at this stage we are unable to mechanistically define the link between CLN7 dysfunction and these cellular defects, we present compelling evidence that this neuropathology has a significant nuclear component that does not present in the *Cln7^Δex2^* mouse and is most likely caused by gain-of-function of aberrant CLN7 protein. There is emerging evidence for gain-of-function caused by CLN7 mutation in other neurodegenerative spectrum diseases where CLN7/MFSD8 mutation has been shown to contribute to ALS/FTD (*20*, *36*). Some allelic mutations in CLN7 can contribute to retinopathies including retinitis pigmentosa and macular dystrophy (*37*–*39*). CLN7 has been shown to co-localize with post-synaptic density marker 95 (PSD-95) in the mouse retina indicating that it may play more complex roles in specialist neural cell-types (*39*). There is an interesting parallel where TDP-43 mutation in ALS/FTD results in defective nuclear pore complex function (*40*, *41*); CLN7 could act by a similar mechanism. In these variant CLN7 diseases, the missense mutation in a single allele may exert a dominant negative effect that, when combined with other genetic or environmental factors, results in the manifestation of disease. Therefore, the known phenotypes of CLN7 disease from adult retinal dystrophy to late infantile disease may actually represent a combination of a dosage-responsive disease spectrum proportional to mutation severity and an element of gain-of-function that is mutation-specific.

The assumption that CLN7 exists as a single 518 aa protein of 57kDa is not consistent with Ensembl predictions of up to 24 protein coding transcript variants and our extensive western blot evaluations showing up to 12 discrete bands ranging from 33-180kDa in multiple cell types. The larger molecular weight bands were unchanged under non-reducing conditions or after PNGase F digestion although this did reduce a 60kDa band to 57kDa in SH-SY5Y proving that the full-length CLN7 form is subject to N-glycosylation. How this relates to function remains to be elucidated. We hypothesize that the isoforms smaller than 57kDa are caused by alternate mRNA splicing and/or post-translational modifications including protease cleavage. Isoforms consistent across HEK293T, SH-SY5Y, MEFs and iNPC included western bands at 57, 50, 44, 40, 37 and 33kDa that equated to predicted sizes for putative proteins encoded by transcript variants with and without Cys^401^ protease digestion (**Table 3**). The flCLN7 57kDa protein is reduced to 44kDa by Cys^401^ digestion and a transcript variant CLN7^Δex7/8^ 50kDa protein is reduced to 37kDa by Cys^401^ digestion. Nuclear fractionations in iNPC reveal that the 50kDa CLN7 isoform localizes to the nucleus in control and BD iNPC but the 37kDa is only present in BD iNPC. We generated plasmids expressing the 57, 50, 44 and 37kDa isoforms designed by recapitulating Δex7/8 and Cys^401^ cleavage in CLN7 coding sequence and transfected *Cln7^Δex2^* knockout MEFs. As expected, flCLN7 and CLN7^Cys401^ show vesicular staining that partially overlaps with p62, and CLN7^Δex7/8^ and CLN7^Δex7/8Cys401^ show substantial localization to the nucleus. This was further confirmed by transfecting post-mitotic *Cln7^Δex2^* mouse cortical neurons with the BD specific CLN7^Δex7/8Cys401^ 37kDa isoform. Finally, we sought to evaluate whether over-expression of these CLN7 isoforms induced apoptosis. Transfection of all three truncated isoforms causes elevated apoptosis in HEK293T cells compared to a GFP expressing control but the CLN7^Cys401^ transgene results in a 7-fold increase in apoptosis induction. This infers that flCLN7 protease cleavage at vesicular membranes can produce a pro-apoptotic protein isoform. This is consistent with apoptosis being linked with bioenergetically impaired mitochondria and cytochrome c release and could be a possible mechanism for decreased gene therapy efficacy on dose escalation.

Overall, our study shows that gene therapy using a full-length transgene encoding the 518aa form of CLN7 driven by a strong neuronal promoter ameliorates some but not all aspects of BD phenotype in the *Cln7^Δex2^* knockout mouse. Moreover, dose escalation shows concerning signs that high levels of CLN7 expression could have a negative impact on transduced neurons. This led us to investigate the nature of CLN7 in human disease. Using iPSC-derived iNPC we have shown that CLN7 disease patients, where disease is caused by two different missense mutations, do express multiple different isoforms of CLN7 that aberrantly localize within iNPC thereby probably contributing to disease. Such disease mechanisms cannot occur in the knockout mouse model which does not express CLN7. We identified a 37kDa CLN7 isoform, consistent with exon 7/8 splice excision and subsequent protease digestion at Cys^401^, that accumulates in the nuclei of CLN7 disease patient iNPC. We predict the mechanism of disease pathology by this isoform is through a gain-of-function effect. CLN7 iNPC present with a profound deficit of the nuclear proteome, particularly of the nuclear pore complex, so it is plausible that the 37kDa isoform impairs nuclear transport. CLN7 iNPC and *Cln7^Δex2^* cortical neurons share mitochondrial bioenergetic defects which relate to loss of protein function, although this could relate to flCLN7 or the protease cleavage CLN7^Cys401^ isoform. Critically, when expressed at high levels, CLN7^Cys401^ causes apoptosis, likely through mitochondrial cytochrome c release. Minimal Essential Network analysis of our proteomics data identifies dominator proteins involved in mitochondrial quality control which could infer a role for this cleavage isoform as well as explaining the accumulation of SCMAS in lipofuscin characteristic of Batten disease. In conclusion, it is plausible that CLN7 dose escalation could result in therapy and toxicity in equal measure. This has implications in the design of optimized gene therapy for CLN7 disease and in future clinical trials. Further, this could be considered a wider paradigm for genetic medicines where transgenic protein isoforms and dosage may play significant roles.

## Materials and Methods

### AAV plasmid construction and viral vector production

The human cDNA sequence for full length *CLN7* (1554nt, 518aa encoding a predicted 57kDa protein) was cloned into a single-stranded AAV9 expression construct (**Supp Fig 1**). Gene expression was driven by the human *synapsin 1* promoter and the construct included a downstream woodchuck hepatitis virus regulatory element (WPRE). An equivalent control construct carrying eGFP was also produced. AAV virus was produced by triple transfection of HEK 293T cells with harvesting of viral particles after 72h via iodixanol gradient purification and ultracentrifuge. AAV particles were concentrated using a Vivaspin 20 centrifugal concentrator with a 100,000 mw cut-off (Sartorius Stedim Biotech, UK). This method has been previously described (*42*). The verification of successful virus production was confirmed via SDS-Page to visualize the presence of capsid proteins and TaqMan PCR targeting the synapsin promoter sequence (F: TGCCTACCTGACGACCGAC R: GGTGCTGAAGCTGGCAGTG) to determine the viral genome titre.

### Animal procedures

All animal experiments were performed in following approval by local research ethics governance and in accordance with the Animal Scientific Procedures Act 1986, UK or European Union Directive 86/609/EEC and Recommendation 2007/526/EC, regarding the protection of animals used for experimental and other scientific purposes. Procedures were conducted at UCL under the Home Office Project Licence PCC436823, following ARRIVE guidelines and recommendations. Animals were bred in cages (maximum of five animals per cage), and a light–dark cycle was maintained for 12h. The humidity was 45–65%, and the temperature was 20–25 °C. Animals were fed *ad libitum* with a standard solid diet (17% proteins, 3% lipids, 58.7% carbohydrates, 4.3% cellulose, 5% minerals and 12% humidity) and given free access to water. Cln7 knockout mouse carrying the European Conditional Mouse Mutagenesis (EUCOMM) tm1d allele by *Cre*-mediated recombination of the floxed exon 2 of the murine *Cln7/Mfsd8* gene (*Cln7^Δex2^*)(*18*) were used.

### Mouse genotyping

For Cln7^Δex2^ genotyping, a PCR with the following primers was performed: TGGTGCATTAATACAGTCCTAGAATCCAGG, CTAGGGAGGTTCAGATAGTAGAACCC, TTCCACCTAGAGAATGGAGCGAGATAG-3’, resulting in a 290 bp band in the case of Cln7^Δex2^ mice, and 400 bp for wild-type ^(*18*)^.

### *In vivo* gene therapy

Congenic wild-type and *Cln7^Δex2^* mice (*18*) were bred as homozygous colonies on a C57Bl/6 background. Newborn P0-P1 mouse pups (n=40) received 2.5×10^11^ vg (high dose) of AAV9.SYN1.CLN7 vector or 2.5×10^10^ vg (low dose) of either AAV9.SYN1.CLN7 or AAV9.SYN1.eGFP vector via bilateral intracerebroventricular injections using a 3 3-gauge needle (Hamilton, USA) to each lateral ventricle, as described previously (*43*). After injection, the pups were returned to the dam and their weight was monitored daily for the first week. Mouse weights were recorded monthly.

### Animal behavioural analysis

The wild-type, untreated and treated *Cln7^Δex2^* mice were subjected to a range of behavioral tests to examine for differences in the performance of the different treatment groups.

### Accelerating rotarod test

Mice were placed on the accelerating rotarod (Harvard Apparatus, MA) programmed to increase from an initial rotation speed of 4-45 rpm over a 5-minute period. The mice underwent training before as described (*19*) and were assessed monthly with two trials on the same day and with a rest period between. The time taken for the mouse to fall from the rod was recorded and used as a measure of performance.

### Foot-fault test

The mice were placed on a metal grid (35×40cm) with a mesh size of (1.3×1.3cm) suspended above the bench. The mouse was allowed to explore the grid for 1 minute and was filmed over this period. The footage was then analysed with a foot-fault being recorded when the foot passed through the grid. The percentage of missteps out of total steps was then calculated.

### Open field test

The test mice were placed into a white box (48×48cm) and allowed to acclimatise to their new surroundings for 1 minute before being recorded for 4 minutes. The footage was then analysed using the *ANY-maze* software (Anymaze, Dublin, IR) to measure various parameters such as average speed, distance travelled, time in the centre of the box, entries from periphery to center, rotations and time stationary.

### Vertical pole test

Mice were placed at the top of a wooden pole (diameter - 1.4cm, height-28cm) facing downwards and their descent of the pole was observed. If the mouse was able to descend the pole in a controlled manner within a 1-minute window, the test was considered successful and the mouse scored 1 for a pass otherwise it was considered a fail and the mouse scored 0.

### Brain tissue sectioning

Upon reaching the end point of the study, either based on the humane endpoint or pre-determined at 9 and 14-months, the mice were collected via terminal exsanguination by trans-cardiac perfusion with phosphate buffered saline. The organs were extracted from the mice and fixed in 4% paraformaldehyde for 48 hours and transferred to a 30% sucrose in phosphate-buffered saline for cryoprotection. Subsequently the brains were sectioned using a cryostat at −20°C to a thickness of 40μm in the coronal plane.

### Tissue immunohistochemistry

Brain sections were stained for marker proteins for microglia (CD68) and Astrocytes (GFAP), lysosomal membrane protein LAMP1 and neurons (NeuN). Free-floating brain sections were incubated in a 1% H_2_O_2_ solution for 30 minutes to block endogenous peroxidase activity followed by three washes with TBS. A block of non-specific binding was then performed by a 30-minute incubation in 15% normal serum with 0.3% TBS-T. The solution was then changed to a 10% normal serum in 0.3% TBS-T with the primary antibodies added (anti-CD68, Bio-Rad MCA1957; anti-GFAP, Millipore MAB377; anti LAMP1, ab24170 Abcam, anti-NeuN, MAB377 Abcam, anti-MFSD8 PA5-60832 ThermoFisher) and incubated overnight at 4°C. The sections were washed 3 times with TBS followed by incubation in biotinylated secondary antibody (anti-rabbit, anti-rat or anti-mouse, 1:1000, Vector Laboratories) in 10% normal serum with 0.3% TBS-T for 2 hours. After three further washes with TBS, visualization of the staining was achieved with Vectastain avidin-biotin solution (Vector Laboratories) and DAB (Sigma). The sections were then mounted, dehydrated, cleaned in histoclear and coverslipped with DPX (VWR).

For visualization of autoflourescent storage material, the sections were firstly mounted onto Superfrost Plus slides and allowed to air dry for 20-minutes before coverslipping using Flouromount-G with DAPI mounting medium (Southern Biotech).

### Thresholding image analysis and cortical thickness measurements

To analyze the degree of autofluorescent storage material accumulation (AFSM) and glial activation (GFAP-positive astrocytes and CD68-positive microglia), a semi-automated thresholding image analysis was used with Image-Pro Premier software (Media Cybernetics, MD). Briefly, slide-scanned images at 10x magnification for stained sections were collected for all animals using a Zeiss Axioscan Z1 (Zeiss Microscopy Deutschland GmbH followed by demarcation of anatomical regions of interest. Images were subsequently analyzed using Image-Pro Premier (Media Cybernetics) using appropriate thresholds that selected the foreground immunoreactivity above background. Separate thresholds were used for each antigen and each anatomical region analyzed. Cortical thickness was measured in brain sections stained with Anti-NeuN antibody using *StereoInvestigator* software (MBF Bioscience, VT). This was done by drawing 10 individual contours over the entire thickness of cortex across 3 separate sections and collecting the average value in μm.

### Mouse embryonic fibroblast (MEF) isolation and culture

MEFs were prepared from fetal (E13.5) *Cln7^Δex2^* pregnant (*18*) or +/+ pregnant (WT) mice according to a previously validated protocol (*44*). Briefly, each embryo was dissected into 1 ml sterile Eagle’s Basal Salt Solution (EBBS), voided of its internal organs, and incubated with 1 ml of 0.25% trypsin-EDTA for 20 minutes at 37°C. The tissue was triturated and the cell suspension seeded on 60 cm^2^ Petri dishes in high glucose (25□mM) DMEM (Sigma, Madrid, Spain) with 10% fetal calf serum (Roche Diagnostics, Heidelberg, Germany) and L-glutamine (4□mM), and incubated at 37°C in a humidified 5% CO_2_-containing atmosphere.

### Primary cortical neuron isolation and culture

Primary cultures of mice cortical neurons were prepared from the offspring of 14.5 days *Cln7^Δex2^* pregnant (*18*) or +/+ pregnant (WT) mice according to a previously validated protocol (*44*). In essence, cells were seeded at 2.0×10^5^ cells per cm^2^ in different-sized plastic plates coated with poly-D-lysine (10 μg/mL) and incubated in Neurobasal-A (Life Technologies) supplemented with 5.5 mM of glucose, 0.25 mM of sodium pyruvate, 2 mM glutamine and 2% (vol/vol) B27 supplement (Life Technologies). At 72 h after plating, medium was replaced, and cells were used at day 7. Cells were incubated at 37 °C in a humidified 5% (vol/vol) CO_2_-containing atmosphere.

### Cell line culture

HEK293T cells were cultured in complete Dulbecco’s Modified Eagle Medium (DMEM) (Gibco^™^) (10% Fetal Bovine Serum, 2% L-Glutamine, 1% Penicillin/Streptomycin and 0.2% MycoZap^™^). SH-SY5Y cells were cultured on complete DMEM/F-12 (Gibco^™^) (10% Fetal Bovine Serum, 2% L-Glutamine, 1% Penicillin/Streptomycin and 0.2% MycoZap^™^).

### Culture of human hiPSC

iPSC lines were previously generated as described using episomal doggybone DNA vectors (*45*) using dermal fibroblasts held by the UCL NCL database and cell repository (https://www.ucl.ac.uk/ncl-disease/), donated by two CLN7 patients (Pa380; homozygous for c.881C>A, p.(Thr294Lys) and Pa474; homozygous for c.1393C>T, p.(Arg465Trp)) and two age-matched controls (PromoCell). All iPSC were maintained on inactivated MEFs (iMEFs) feeder layer and transitioned to feeder-free in preparation for neural specification. iPSC maintenance medium is composed of DMEM/F-12 (Gibco^™^), 2% L-Glutamine, 20% Knockout^™^ Serum Replacement (Gibco^™^), 1× MEM Non-Essential Amino Acids Solution (100×) (Gibco^™^), 1× N-2 Supplement (100×) (Gibco^™^), 0.1 mM β-mercaptoethanol (50 mM), 10 ng/ml Recombinant Human FGF-basic (PrepoTech^®^), 1% Pen/Strep and 0.2% MycoZap^™^ Prophylactic (Lonza). Essential 8^™^ Medium (Gibco^™^) was prepared for feeder-free iPSCs culture on Matrigel^®^ Matrix by addition of the Essential 8^™^ Supplement (50 ×) to the 500 mL Essential 8^™^ Basal Medium according to the manufacturer’s instructions, 1% Pen/Strep and 0.2% MycoZap^™^ Prophylactic (Lonza).

### Neural specification of iPSC

Plates and flasks for feeder-free iPSCs, NSCs and NPCs culture were previously coated with Matrigel^®^ Matrix (Corning^®^). For iPSCs culture on feeders, iMEFs were plated on previously coated wells with autoclaved 0.1% gelatin. Neural Induction Medium (NIM) was prepared and used for culturing neurospheres and NSCs, and it is composed of DMEM/F-12 (Gibco ^™^), 1× MEM Non-Essential Amino Acids Solution (100×) (Gibco^™^), 1× N-2 Supplement (100×) (Gibco^™^), 20 ng/ml Recombinant Human FGF-basic, 0.1% Heparine, 1 % Pen/Strep and 0.2% MycoZap^™^ Prophylactic (Lonza). Neural Expansion Medium (NEM) was employed for culturing NPCs, and it is formed of DMEM/F-12 (Gibco ^™^), 1× MEM Non-Essential Amino Acids Solution (100×) (Gibco^™^), 1× N-2 Supplement (100×) (Gibco^™^), 1× B-27^™^ Supplement (50×) (Gibco ^™^), 20 ng/ml Recombinant Human FGF-basic, 0.1% Heparine, 1% Pen/Strep and 0.2% MycoZap^™^ Prophylactic (Lonza).

iPSCs colonies were cultured on Matrigel^®^ Matrix on 6-well plates at 80% confluency. Cells were washed with PBS and dissociated with TrypLE^™^ Express (Gibco^™^). They were centrifuged at 107 × g for 8 min and resuspended in Essential 8^™^ supplemented medium. Cells were counted using a Neubauer chamber and 7.5 × 10^3^ cells per well were plated on V-Shaped Bottom plates (low-attachment 96-well plates) in 100 μl of Neural Induction Medium (NIM) and ROCK inhibitor (Y-27632 dihydrochloride, 1254, Biotechne^®^) (used at 1:1000). Half of the medium (50 μl) was changed every day during 5 days after cell plating to allow cells to form spheroids. After 5 days of medium change, 10-15 spheroids per well were plated in 6-well plates previously coated with Matrigel^®^ Matrix (Corning^®^). NSCs (neural rosettes and neural tubes) were dissociated with TrypLE^™^ Express and centrifuged at 107 × g for 8 min for neural differentiation. The cell pellet was resuspended in Neural Expansion Medium (NEM) and cells were plated on new 6-well plates coated with Matrigel^®^ Matrix to generate NPCs which grow as single cells.

### Plasmids and transfections

Plasmid; pEGFPC1-CLN7, pCMV10-3xFLAG-CLN7 were a kind gift from Dr. Stephan Storch (University of Hamburg) and have been previously described Steenhuis *et al.* (*30*) The following plasmids contain CLN7 transgene variants and were *de novo* synthesized by GeneArt (Thermo Fisher); pcDNA3.4-c1CLN7/8, pcDNA3.4-e7/8CLN7, pcDNA3.4-e78CLN7-C1 (**Supplementary Table 1**). iNPC, MEFs, primary cortical neurons, 293T and SH-SY5Y cells were transfected using Lipofectamine LTX reagent (Thermo Fisher) according to the manufacturer’s protocol with 2 μg DNA per 9.6 cm^2^ well and cells analyzed 24-72 hours later.

### Bafilomycin A treatment

iNPC and primary cortical neurons were treated for 6 hours with 100 nM bafilomycin A (1334, Tocris) for the bioenergetics and immunocytochemistry analyses.

### Bioenergetics analysis

Oxygen consumption rate (OCR) of iNPCs and *Cln7^Δex2^* mouse cortical neurons were measured in real-time in an XFe24 Extracellular Flux Analyzer (Seahorse Bioscience; Seahorse Wave Desktop software 2.6.1.56). The instrument measures the extracellular flux changes of oxygen in the medium surrounding the cells seeded in XFe24-well plates. Assays wer performed 1 to 7 days after cell plating. Regular cell medium was removed and cells were washed twice with DMEM running medium (XF assay modified supplemented with 5 mM glucose, 2 mM L-glutamine, 1 mM sodium pyruvate, 5 mM HEPES, pH 7.4) and incubated at 37°C without CO_2_ for 30 minutes to allow cells to pre-equilibrate with the assay medium. Oligomycin, FCCP or antimycin/rotenone diluted in DMEM running medium were loaded into port-A, port-B or port-C, respectively. Final concentrations in XFe24 cell culture microplates were 1 μM oligomycin, 2 μM FCCP and 2.5 μM antimycin and 1.25 μM rotenone. The sequence of measurements was as follows, unless otherwise described. Basal level of OCR was acquired in 3 consecutive determinations, and thereafter oligomycin was injected and mixed for 3 minutes before the OCR was acquired in 3 consecutive determinations. The same procedure was followed for FCCP and antimycin/rotenone. All determinations were normalized to the basal OCR obtained in the corresponding well. Three to five replicas were determined for each sample obtained from the number of independent cell culture preparations indicated in the figure legends. The non-mitochondrial OCR was obtained after antimycin/rotenone injection, and the maximal respiration was obtained after FCCP injection minus that of the non-mitochondrial OCR. ATP production was calculated by subtracting, to the last OCR value before oligomycin injection, that obtained after oligomycin injection.

### Cell immunocytochemistry

iPSCs and iNPCs cultured on 12-well plates or on Nunc Lab Tek chamber slides (8 wells) were washed twice with PBS and fixed with 100% ice-cold methanol for 5 min (on ice or at −20□C). Cells were washed again with PBS three times. Cells were blocked with blocking solution (2% BSA in PBS, 1% goat serum, 0.1% Triton X-100) for 30 min on the rocker. Primary antibodies anti-TRA-1-60 (ab16288, Abcam) and anti Oct3/4 (AF1759, R&D Systems) were prepared in the blocking solution at 1:200 dilution; and anti-SOX2 (AF2018, R&D Systems), anti-Nanog (ab80892, Abcam) and anti-MFSD8 (PA5-60832, Thermo Fisher) were also prepared in the blocking solution at a 1:100 dilution. Cells were incubated with primary antibodies for 2 h at room temperature and washed three times with PBS. Secondary antibodies (A110088, Alexa Fluor ^™^ 488 goat anti-rabbit; A11055, Alexa Fluor ^™^ 488 donkey anti-goat; A11004, Alexa Fluor^™^ 568 goat anti-mouse; and A11011, Alexa Fluor ^™^ 568 goat anti-rabbit) were prepared at 1:500 dilution in blocking solution and cells were incubated with them for 1 h at room temperature. Cells were washed three times with PBS. To stain cell nuclei, 1 mg/ml DAPI solution (D9542, Sigma-Aldrich) used at 1:1000 in PBS was added to the cells for 15-30 min or a drop of the DAPI Mounting (Vectashield^®^) solution was added to each well. In the first case, cells were washed three times with PBS, and they were ready to image after the addition of PBS to each well. In the second case, cells were incubated at 4□C overnight before imaging them. Cells were imaged using the Leica 6000CTR Live cell imaging fluorescent microscope (for plates) and the Zeiss Imager M2 microscope (for slides).

MEFs and cortical neurons were seeded on coverslips, fixed with a 4% paraformaldehyde (PFA) solution, blocked, and incubated with primary antibodies overnight at 4°C. The primary antibodies were anti-GFP (1/500) (ab655; Abcam), anti-MFSD8, (1/100) (PA5-60832; Invitrogen), anti-LAMP1 (1/100) (1D4B; Developmental Studies Hybridoma Bank) and anti-p62 (1/100) (Abnova clone 2c11-H00008878-M01). They were then incubated for 1 h with fluorescent secondary antibodies (1/500) Alexa Fluor 568 antimouse (A-10037; Thermo Fisher), Alexa Fluor 488 anti-rabbit (A-11008; Thermo Fisher), and Alexa Fluor anti-rat 568 (A-21247; Thermo Fisher). DAPI (4’,6-diamidino-2-phenylindole) was used for nuclei visualization. Coverslips were mounted in ProLong Gold antifade reagent. Negative controls were performed with either no primary or no secondary antibodies. No staining was detected in any case. Images were acquired on a Dragonfly 200 (ANDOR) imaging system using 100x (1.45 numerical aperture) objective. Images were acquired at the same exposure times in the same imaging session using Fusion 2.2 (NEW) software. Quantification was performed after appropriate thresholding using ImageJ software (NIH) in a minimum of 30 cells from at least 3 experiments. Colocalization between the anti-MFSD8 staining (green) and DAPI was used as a nuclear CLN7 indicator. Percentage was calculated using the number of positive cells for colocalization between total ones. Mitotic cells were defined using DAPI staining. Cells with condensed chromatin and the loss of nuclear membrane were considered as a mitotic cell. Stages of mitoses that were included were representative of prophase through anaphase. For fluorescence confocal microscopy, cells were fixed in 100% ice-cold methanol. Samples were blocked in 2% BSA in PBS +0.05% (v/v) Tween 20 (all from Sigma). Antibodies were diluted in blocking buffer (Antibodies used: MFSD8 1:100 (PA5-60832/ Invitrogen), Nestin 1:200 (Ab18102/ Abcam), Lamp1 1:200 (Ab24170/ Abcam), P62 1:200 (Ab56416)) and incubated overnight at 4 C°. Secondary anti-bodies (1:500, all Alexa Fluor) were applied for 1 hr at room temperature. Immuno-stained cells were mounted with DAPI and analysed using the Leica TCS SP5 X microscope.

### Nuclear fractionation

SH-SY5Y and iNPCs were harvested and centrifuged at 190 × g for 5 min. Cell pellet was washed with PBS and span down at 190 × g for 2 min. A cytoplasmic homogenisation buffer (10 mM HEPES pH 7.9, 60 mM KCl, 1 mM EDTA, 1 mM PMSF, 0.075% v/v NP-40 and PIC) was added to the pellet, vigorously vortexed, incubated on ice for 4 min and vortexed again. This mix was centrifuged at 1500 × g at 4□C for 10 min - the supernatant is the cytoplasmic fraction. The pellet was resuspended in the nuclear extraction buffer (20 mM Tris-HCl, 420 mM NaCl, 1.5 mM MgCl_2_, 0.2 mM EDTA pH 8, 1 mM PMSF, 25 % v/v glycerol and PIC) and incubated on ice for 30 min homogenising it with a syringe (25 G) 10 times. The mix was centrifuged at 17000 × g and 4□C for 40 min – being the supernatant the nuclear fraction. The cytoplasmic fraction was detected by western blotting using anti-ERK1/2 as a marker for the cytoplasmic fraction and anti-Lamin B1 as a marker of the nuclear fraction.

### PNGase F treatment

PNGase F (9109-GH, R&D Systems) was used to remove any possible N-linked glycans from SH-SY5Y cell lysates. Prior to PNGase F treatment, sample denaturation was conducted in a 30 μl reaction volume using 1x Denaturing buffer (10x Denaturing buffer composition: 5% SDS and 50 mM DTT) and incubated at 100 DC for 10 min. The reaction mixture was cooled to room temperature and microcentrifuged. Then 3 μl of 10% Triton^®^ X-100 were added to a final concentration of 1.67%. PNGase F. PNGase F was diluted to 0.167 ng/μl. The reaction mixture was divided in two and 15 μl were combined with 15 μl 0.167 ng/μl and the other 15 μl of reaction were mixed with 15 μl of Assay Buffer (0.1 M Tris, pH 7.5) as a control. The PNGase reaction mixtures were incubated at 37 □C for 30 min followed by the procedure of immunoblotting.

### Western blot

After cell harvesting, equal amounts of protein (measured by BCA assay) were mixed with 5× SDS-loading buffer, incubated for 10 min at 95.5 □C, separated by SDS-PAGE (10% polyacrylamide separating gel and 4% polyacrylamide stacking gel) and blotted onto a PVDF membrane (Millipore) for 2 h at 100 V using a tank transfer. After membrane blocking with 5% BSA in 1× TBST buffer, western blots were probed with the primary antibodies anti-MFSD8 (PA5-60832, Thermo Fisher) and anti-β-actin (A2228, Sigma-Aldrich) at 4°C overnight, followed by detection with HRP secondary antibodies (Polyclonal Goat Anti-Mouse, PO447 Dako, at 1:2000; and Stabilized Peroxidase Conjugated Goat Anti-Rabbit, 32460 ThermoFisher, at 1:500) and visualized using Immobilon Western Chemiluminescent HRP Substrate (WBKLS0500, Millipore) and LI-COR Odyssey Fc. The primary antibody anti-MFSD8 prepared in the blocking solution at 1:500 dilution; and the anti-ß-actin primary antibody was prepared at 1:10000 dilution. MFSD8 protein band intensities were measured and compared to the total MFSD8 protein on each cellular fraction using ImageJ. To probe blotting membranes with different antibodies of similar molecular weight, blotting membranes were stripped using a 0.5 M NaOH solution for 20 min at room temperature on the rocker.

### Flow cytometry

HEK293T cells and NPCs were harvested and washed with PBS. Cells were incubated with eBioscience^™^ Fixable Viability Dye eFluor^®^ 780 (65-0865, ThermoFisher) at 1 μL/mL of cells in azide-free and serum/protein-free PBS for 20-30 min at 4□C. Cells were washed with PBS and centrifuged at 300 × g for 5 min. Cell pellet was resuspended in 100 μL of cell staining solution (2% BSA in PBS) and incubated with 400 nM Apotracker^™^ Green (427402, BioLegend^®^) (diluted in staining solution) for 20 min at room temperature. Another cell wash with PBS was conducted and cells were fixed with cell fixation buffer (4% PFA) for 15-20 min at room temperature. Cells were washed with PBS and 200-500 μL of PBS was added to the cell pellets for flow cytometry analysis in Falcon^®^ 5 ml Round-Bottom Tubes. A MACSQuant from Miltenyi Biotec was used as a flow cytometer. Apotracker^™^ Green was excited using the blue 488 nm laser, and the fluorescence emission was collected using a 525/50 nm laser. The eBioscience^™^ Fixable Viability Dye eFluor^®^ 780 was excited using the red 638 nm laser, and the fluorescence emission was collected using a 785/62 nm laser. The obtained data were analyzed using FlowLogic software.

### Statistical analyses

All data are presented as means ± SEM. n numbers, statistical tests and p values for each experiment are detailed in the respective figure legends. All statistical analysis was performed using GraphPad Prism (GraphPad Software, San Diego, California USA). Unless otherwise stated the following convention is used to denote significance - * p≤0.05, **≤0.01, ***≤0.001 and ****≤0.0001.

### Proteomic analysis

#### Cell Lysis

75 cm^2^ Flasks were used to culture cells to sub-confluent cultures (around 6 million cells per flask). NPCs were washed with PBS then lysed using 0.5 ml lysis buffer composed of 1M TEAB (Triethylammonium bicarbonate) and 0.1% (w/v) SDS (Sodium dodecyl sulphate) for 5 Minutes. Lysates were collected using cell scrapers and aliquots were frozen at −80°C.

#### iTRAQ labelling

The method was performed following the published protocols by Unwin, *et al* (*46*). Protein concentration was determined by Bradford assay (Bio-RAD). Subsequently, 100μg of protein from each sample was equalized to 55μl using 1M TEAB. 0.1 volumes of 50 mM Dithiothreitol (DTT, Fluka) was added to reduce disulphide bonds and incubated at 60°C for an hour. Samples were alkylated by 0.05 volumes of 200mM Iodoacetamide (IAA) at room temperature for 10 minutes. Proteins were digested using 10 μg trypsin (Sequencing Grade Modified Trypsin, Promega) for 100μg of protein in each sample. Trypsin was reconstituted in 1M TEAB so that final SDS concentration in the samples is less than 0.05%, trypsin added to samples and incubated at 37°C overnight. After digestion samples were dried using a SpeedVac concentrator (Labconco) and frozen at −20°C.

Dried protein digests were resuspended using 1 M TEAB to 30 μl. iTRAQ reagent vials (AB Sciex) were resuspended with 70 μl isopropanol (Fluka) and transferred to a protein digest. All samples were allowed to react at room temperature for two hours. Sample volumes were reduced to 30 μl using SpeedVac concentrator to remove isopropanol and stored at −20°C.

#### High pH Reverse Phase Liquid Chromatography Fractionation

Fractionation as performed as described in details in (*47*). Peptides were fractionated off-line using an Agilent 1200 series LC system loading 900ul of sample onto a high pH reversed-phase chromatography column (Agilent ZORBAX 300 Extend-C18 4.6×150mm 3.5 micron). Samples were eluted using a solvent gradient from 5% buffer B and increasing to 30% (buffer B in 35 min, and then to 45% buffer B after 1 min at 45°C and a flow rate of 700 μl/min. Fractions were collected in a 96-well plate, dried using SpeedVac concentrator and stored at −20°C.

#### Strong Anion Exchange Desalting and Low pH Reverse Phase Liquid Chromatography and Mass Spectrometry (LPLC-MS/MS)

Samples were desalted using POROS 50 HQ strong anion exchange beads (1255911, Thermo Fisher) as previously described in detail (*48*). Collected samples were dried completely using SpeedVac concentrator and samples were stored at −20°C. Before Mass Spectrometry, the wells in the plate with samples were thawed and each well that contained samples was resuspended in 10μL of 5% (v/v) acetonitrile / 0.1% (v/v) formic Acid. Samples were then transferred to a sample vial and placed on the sample rack for analysis. Low-pH reversed-phase liquid chromatography using a nanoACQUITY UHPLC system (Waters) and BEH C18 Column (1.7 μm, 75 μm x 250 mm) and separated over a 90 min solvent gradient from 3% ACN (v/v) to 40% ACN (v/v) at a flow rate of 300 nl/min and column temperature at 30 °C. Peptides were eluted on-line to TripleTOF^®^ 6600 Quadrupole Time-Of-Flight (QTOF) Mass Analyzer (AB SCIEX). Spectra were collected using a standard IDA method with a 0.1 second MS followed by 10 MS/MS scans over the next one second with iTRAQ collision energies enabled. Each ion was selected a maximum of two times and was dynamically excluded (± 50 mmu) for 90 seconds thereafter.

#### Data Analysis

After acquisition of fractioned peptides, raw data files were uploaded to Protein Pilot analysis software to identify and quantify peptides *in silico* using ProteinPilot version 5.0.1, AB SCIEX. Proteins were searched against a human-specific Uniprot database (Swiss-Prot Human database, containing 20,371 entries, Version number 10_2019). Detection protein threshold was set to as low as possible. Protein and peptide summaries were exported for statistical assessment. Quantified proteins with unused scores equal or above 2 only were included and false discovery rate was 5%. At least one unique peptide of 99% confidence had to be used to identify the protein (Unused score above 2). Log_2_ for the ratio between samples were used to calculate the fold changes. For the analysis of all the four samples simultaneously across the three mass spectrometry runs identified peptides were coalesced to protein-level quantifications and statistical testing for differential expression performed using v1.0.0 of the in-house developed software ‘BayesProt’ (https://github.com/biospi/bayesprot/releases/tag/v1.0.0). An earlier version of this technique was presented in Freeman *et al* (*49*), which combined Protein-Pilot (AB SCIEX) sample normalization (‘bias correction’) with a hierarchical Bayesian modelling to assess the quantification reliability of each feature and peptide across assays so that only those in consensus influence the resulting protein group quantification strongly. Similarly, unexplained variation in each individual assay is captured, providing both a metric for quality control and automatic down weighting of suspect assays. Each protein group-level quantification were outputted by the software and is accompanied by the standard deviation of its posterior uncertainty. This integrated a flexible differential expression analysis subsystem with false discovery rate control based on the popular MCMCglmm package for Bayesian mixed-effects modelling (MCMCglmm R Package [Hadfield, Jarrod D. “MCMC methods for multi-response generalized linear mixed models: the MCMCglmm R package.” Journal of Statistical Software 33.2 (2010): 1e22.]). Method details are described by Xu, J *et al* in data processing (*50*) and Phillips, A *et al* (*48*).

## Supporting information

Sharaireh et al. bioRxiv 20.04.2022

## Acknowledgements

We thank families and patients for tissue donation and Prof Milan Elleder (Pa380) and Dr Helen Michelakakis (Pa465) for sending clinical detail and cell lines. We acknowledge access to the NCL Resource that lists all known CLN7 mutations and their phenotypic effect in patients. We thank Professor Robin Ali and Dr. Sander Smith (now King’s College London) for providing access to the *Cln7*^Δex2^ mouse colony. We also thank Prof. Mina Ryten, UCL, for help and guidance in navigating GTEx Portal.

T.R.M., A.A.R., J.P.B. and S.E.M. were all recipients of the European Union’s Horizon 2020 research and innovation program under grant agreement no. 666918. A.A.R. is supported by the UK Medical Research Council (MR/N026101/1, MR/R025134/1, MR/S009434/1, MR/S036784/1, and MR/T044853/1), The Sigrid Rausing Trust/UCL Neurogenetics Programme, Wellcome Trust Institutional Strategic Support Fund/UCL Therapeutic Acceleration Support (TAS) Fund (204841/Z/16/Z), Action Medical Research (GN2485) and the NIHR GOSH BRC. S.E.M. acknowledges the support of the Medical Research Council (Award MR/R025134). This work was supported by the National Institute for Health Research Biomedical Research Centre at University College London Great Ormond Street Institute of Child Health and funding to the MRC Laboratory for Molecular Cell Biology University Unit at UCL (award code MC_U12266B) towards lab and office space.

## Author Contributions

Conceptualization; J.P.B., A.A.R., T.R.M. Implementation; A.S., M.G.F., S.H.M., M.G.M., A.P., A.T., M.P.H., O.C.T., H.N., A.E.B., S.F., L.M.F., C.D.T. Data analysis; A.D., R.U. Writing, Reviewing & Editing; A.S., S.H.M., H.N., S.S., S.E.M., J.P.B., A.A.R., T.R.M.

## References

1. S. E. Mole et al., Clinical challenges and future therapeutic approaches for neuronal ceroid lipofuscinosis. Lancet Neurol 18, 107–116 (2019).

2. R. E. Williams, S. E. Mole, New nomenclature and classification scheme for the neuronal ceroid lipofuscinoses. Neurology 79, 183–191 (2012).

3. A. Schulz et al., Study of Intraventricular Cerliponase Alfa for CLN2 Disease. N Engl J Med 378, 1898–1907 (2018).

4. L. Bajaj et al., A CLN6-CLN8 complex recruits lysosomal enzymes at the ER for Golgi transfer. J Clin Invest 130, 4118–4132 (2020).

5. A. di Ronza et al., CLN8 is an endoplasmic reticulum cargo receptor that regulates lysosome biogenesis. Nat Cell Biol 20, 1370–1377 (2018).

6. S. L. Cotman, J. F. Staropoli, The juvenile Batten disease protein, CLN3, and its role in regulating anterograde and retrograde post-Golgi trafficking. Clin Lipidol 7, 79–91 (2012).

7. Y. Wang et al., CLN7 is an organellar chloride channel regulating lysosomal function. Sci Adv 7, eabj9608 (2021).

8. M. E. Bosch et al., Self-Complementary AAV9 Gene Delivery Partially Corrects Pathology Associated with Juvenile Neuronal Ceroid Lipofuscinosis (CLN3). J Neurosci 36, 9669–9682 (2016).

9. J. T. Cain et al., Gene Therapy Corrects Brain and Behavioral Pathologies in CLN6-Batten Disease. Mol Ther 27, 1836–1847 (2019).

10. X. Chen et al., AAV9/MFSD8 gene therapy is effective in preclinical models of neuronal ceroid lipofuscinosis type 7 disease. J Clin Invest, (2022).

11. T. B. Johnson et al., AAV9 Gene Therapy Increases Lifespan and Treats Pathological and Behavioral Abnormalities in a Mouse Model of CLN8-Batten Disease. Mol Ther 29, 162–175 (2021).

12. M. L. Katz et al., AAV gene transfer delays disease onset in a TPP1-deficient canine model of the late infantile form of Batten disease. Sci Transl Med 7, 313ra180 (2015).

13. E. Gardner, S. E. Mole, The Genetic Basis of Phenotypic Heterogeneity in the Neuronal Ceroid Lipofuscinoses. Front Neurol 12, 754045 (2021).

14. J. L. Centa et al., Therapeutic efficacy of antisense oligonucleotides in mouse models of CLN3 Batten disease. Nat Med 26, 1444–1451 (2020).

15. C. J. Minnis et al., Global network analysis in Schizosaccharomyces pombe reveals three distinct consequences of the common 1-kb deletion causing juvenile CLN3 disease. Sci Rep 11, 6332 (2021).

16. C. Soldati et al., Repurposing of tamoxifen ameliorates CLN3 and CLN7 disease phenotype. EMBO Mol Med 13, e13742 (2021).

17. I. Lopez-Fabuel et al., Aberrant upregulation of the glycolytic enzyme PFKFB3 in CLN7 neuronal ceroid lipofuscinosis. Nat Commun 13, 536 (2022).

18. L. Brandenstein, M. Schweizer, J. Sedlacik, J. Fiehler, S. Storch, Lysosomal dysfunction and impaired autophagy in a novel mouse model deficient for the lysosomal membrane protein Cln7. Hum Mol Genet 25, 777–791 (2016).

19. S. M. Kleine Holthaus et al., Neonatal brain-directed gene therapy rescues a mouse model of neurodegenerative CLN6 Batten disease. Hum Mol Genet 28, 3867–3879 (2019).

20. E. G. Geier et al., Rare variants in the neuronal ceroid lipofuscinosis gene MFSD8 are candidate risk factors for frontotemporal dementia. Acta Neuropathol 137, 71–88 (2019).

21. M. Kousi et al., Mutations in CLN7/MFSD8 are a common cause of variant late-infantile neuronal ceroid lipofuscinosis. Brain 132, 810–819 (2009).

22. L. M. FitzPatrick et al., NF-kappaB Activity Initiates Human ESC-Derived Neural Progenitor Cell Differentiation by Inducing a Metabolic Maturation Program. Stem Cell Reports 10, 1766–1781 (2018).

23. C. Mauvezin, T. P. Neufeld, Bafilomycin A1 disrupts autophagic flux by inhibiting both V-ATPase-dependent acidification and Ca-P60A/SERCA-dependent autophagosome-lysosome fusion. Autophagy 11, 1437–1438 (2015).

24. G. Wu, E. Dawson, A. Duong, R. Haw, L. Stein, ReactomeFIViz: a Cytoscape app for pathway and network-based data analysis. F1000Res 3, 146 (2014).

25. H. Yu, P. M. Kim, E. Sprecher, V. Trifonov, M. Gerstein, The Importance of Bottlenecks in Protein Networks: Correlation with Gene Essentiality and Expression Dynamics. PLOS Computational Biology 3, e59 (2007).

26. S. Geisler et al., PINK1/Parkin-mediated mitophagy is dependent on VDAC1 and p62/SQSTM1. Nat Cell Biol 12, 119–131 (2010).

27. A. Zeb et al., A novel role of KEAP1/PGAM5 complex: ROS sensor for inducing mitophagy. Redox Biol 48, 102186 (2021).

28. Q. Zheng et al., Hsp70 participates in PINK1-mediated mitophagy by regulating the stability of PINK1. Neurosci Lett 662, 264–270 (2018).

29. K. K. Deol et al., Proteasome-Bound UCH37/UCHL5 Debranches Ubiquitin Chains to Promote Degradation. Mol Cell 80, 796–809 e799 (2020).

30. P. Steenhuis, J. Froemming, T. Reinheckel, S. Storch, Proteolytic cleavage of the disease-related lysosomal membrane glycoprotein CLN7. Biochim Biophys Acta 1822, 1617–1628 (2012).

31. M. Damme et al., Gene disruption of Mfsd8 in mice provides the first animal model for CLN7 disease. Neurobiol Dis 65, 12–24 (2014).

32. J. K. De Zutter, K. B. Levine, D. Deng, A. Carruthers, Sequence determinants of GLUT1 oligomerization: analysis by homology-scanning mutagenesis. J Biol Chem 288, 20734–20744 (2013).

33. T. Danyukova et al., Loss of CLN7 results in depletion of soluble lysosomal proteins and impaired mTOR reactivation. Hum Mol Genet 27, 1711–1722 (2018).

34. C. Mauvezin, P. Nagy, G. Juhasz, T. P. Neufeld, Autophagosome-lysosome fusion is independent of V-ATPase-mediated acidification. Nat Commun 6, 7007 (2015).

35. D. Lim et al., Ca(2+) handling at the mitochondria-ER contact sites in neurodegeneration. Cell Calcium 98, 102453 (2021).

36. L. Huang et al., Mutation analysis of MFSD8 in an amyotrophic lateral sclerosis cohort from mainland China. Eur J Neurosci 53, 1197–1206 (2021).

37. M. Bauwens et al., Functional characterization of novel MFSD8 pathogenic variants anticipates neurological involvement in juvenile isolated maculopathy. Clinical genetics 97, 426–436 (2020).

38. J. Birtel et al., Next-generation sequencing identifies unexpected genotype-phenotype correlations in patients with retinitis pigmentosa. PloS one 13, e0207958 (2018).

39. K. N. Khan et al., Specific Alleles of CLN7/MFSD8, a Protein That Localizes to Photoreceptor Synaptic Terminals, Cause a Spectrum of Nonsyndromic Retinal Dystrophy. Investigative ophthalmology & visual science 58, 2906–2914 (2017).

40. C. C. Chou et al., TDP-43 pathology disrupts nuclear pore complexes and nucleocytoplasmic transport in ALS/FTD. Nat Neurosci 21, 228–239 (2018).

41. A. N. Coyne et al., Nuclear accumulation of CHMP7 initiates nuclear pore complex injury and subsequent TDP-43 dysfunction in sporadic and familial ALS. Sci Transl Med 13, (2021).

42. M. P. Hughes et al., AAV9 intracerebroventricular gene therapy improves lifespan, locomotor function and pathology in a mouse model of Niemann-Pick type C1 disease. Hum Mol Genet 27, 3079–3098 (2018).

43. J. Y. Kim, S. D. Grunke, Y. Levites, T. E. Golde, J. L. Jankowsky, Intracerebroventricular viral injection of the neonatal mouse brain for persistent and widespread neuronal transduction. J Vis Exp, 51863 (2014).

44. R. Requejo-Aguilar et al., PINK1 deficiency sustains cell proliferation by reprogramming glucose metabolism through HIF1. Nat Commun 5, 4514 (2014).

45. C. D. Thornton et al., Safe and stable generation of induced pluripotent stem cells using doggybone DNA vectors. Mol Ther Methods Clin Dev 23, 348–358 (2021).

46. R. D. Unwin, J. R. Griffiths, A. D. Whetton, Simultaneous analysis of relative protein expression levels across multiple samples using iTRAQ isobaric tags with 2D nano LC-MS/MS. Nat. Protocols 5, 1574–1582 (2010).

47. A. M. Sharaireh, L. M. Fitzpatrick, C. M. Ward, T. R. McKay, R. D. Unwin, Epithelial cadherin regulates transition between the naive and primed pluripotent states in mouse embryonic stem cells. Stem Cells 38, 1292–1306 (2020).

48. A. Phillips, R. D. Unwin, S. Hubbard, A. Dowsey. (Methods in Molecular Biology Springer 2021).

49. O. J. Freeman et al., Metabolic Dysfunction Is Restricted to the Sciatic Nerve in Experimental Diabetic Neuropathy. Diabetes 65, 228–238 (2016).

50. J. Xu et al., Regional protein expression in human Alzheimer’s brain correlates with disease severity. Commun Biol 2, 43 (2019).

