## Supplementary material for "CLN7 mutation causes aberrant redistribution of protein isoforms and contributes to Batten disease pathobiology": Sharaireh et al. bioRxiv 20.04.2022

**Title**

**Mutant CLN7 isoforms contribute to Batten disease**

**Authors**

Aseel M. Sharaireh[1,2,3,4], Marta Guevara-Ferrer[1], Saul Herranz-Martin[5], Marina Garcia-Macia[6,7], Alexander Philips[3,4], Anna Tierney[3,4], Michael P Hughes[5], Oliver Coombe-Tennant[5], Hemanth Nelvagel[5], Alysha E. Burrows[1], Stuart Fielding[1], Lorna M. FitzPatrick[1], Christopher D. Thornton[1], Stephan Storch[9], Sara E. Mole[10], Andrew Dowsey[11,12], Richard Unwin[3,4], Juan P. Bolanos[6,7,8], Ahad A. Rahim[5] and Tristan R. McKay[1]*

**Abstract**

The variant late infantile form of the inherited neurodegenerative Batten disease (BD) is caused by mutations in the CLN7/MFSD8 gene and represents a strong candidate for gene therapy. Post-natal intracerebral administration of AAV9-hCLN7 to *Cln7*$^{\Delta ex2}$ knockout mice resulted in extended lifespan but dose escalation resulted in reduced acuity in neurophysiology tests, cerebral atrophy and elevated neuroinflammation. Comparing patient and control iPSC-derived neural progenitor cells (iNPC) we discovered that CLN7 localizes to the nucleus as well as the endolysosomal network and is differentially distributed in BD iNPC. Proteomics identified a profound nuclear defect in BD iNPC that compounds with mitochondrial and lysosomal metabolic defects resulting in elevated apoptosis. We further identified a 50kDa common nuclear CLN7 isoform and a 37kDa isoform that accumulates only in BD iNPC nuclei. Our findings suggest that successful treatment of CLN7 BD will require combinatorial therapies addressing both loss and aberrant gain of protein function.

**This PDF file includes:**

Supplementary Text
Figs. S1 to S12
Tables S1 to S2

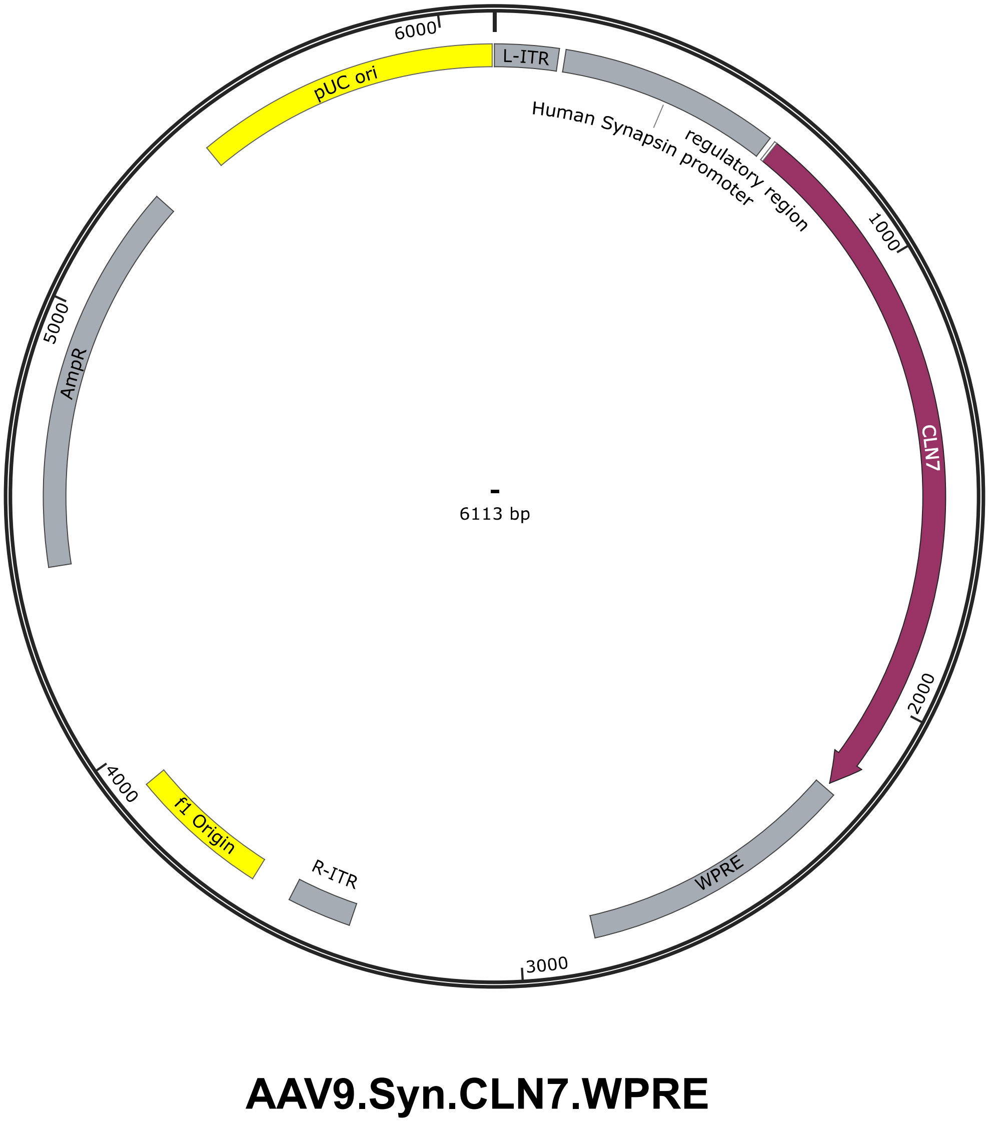


**Fig. S1.** Schematic representation of the pAAV9.Syn.CLN7.WPRE expression vector used for AAV9 gene therapy vector production.

#### WEIGHT

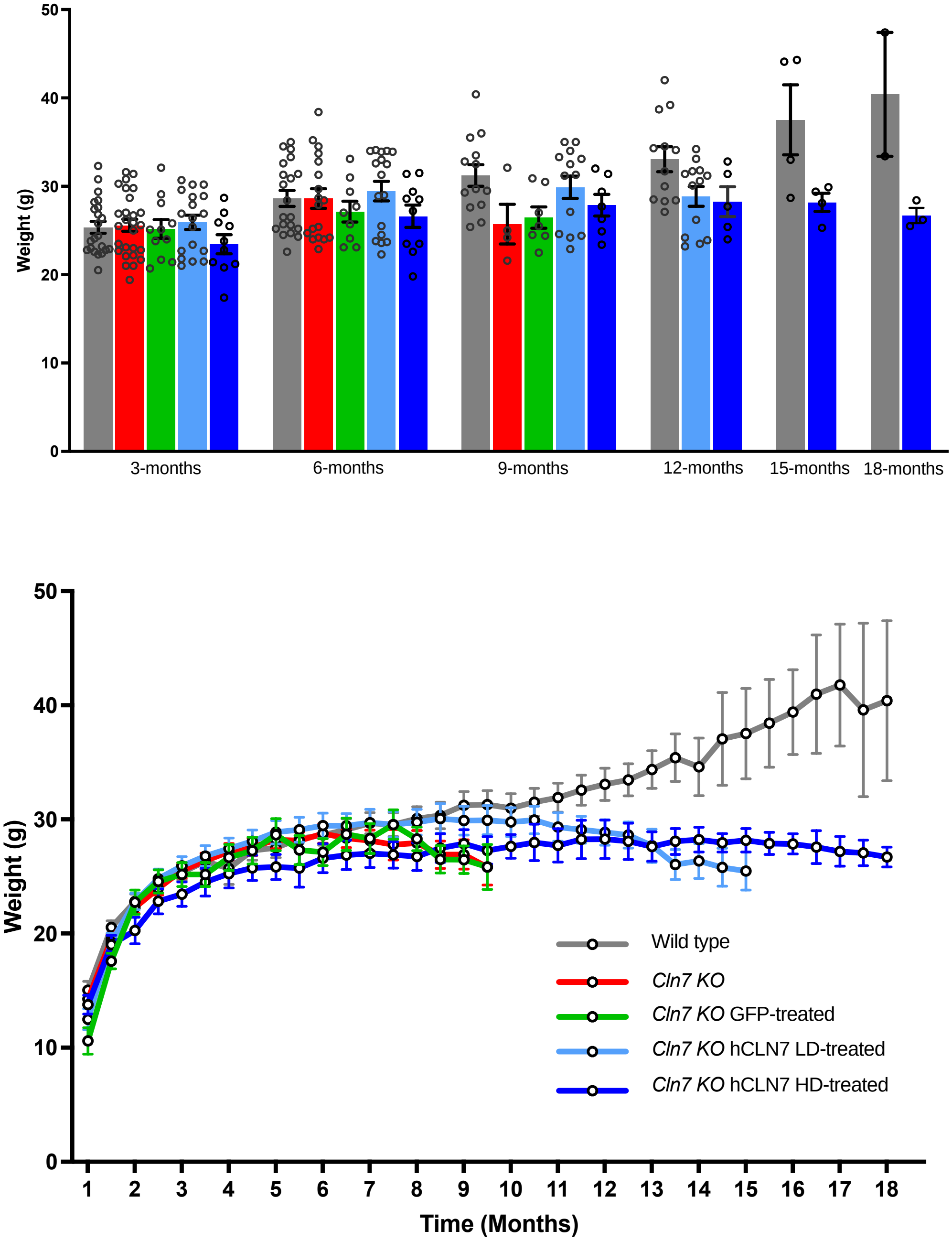


**Fig. S2.** Body weight (g) of wild-type and *Cln7*$^{\Delta ex2}$ mice treated with intracerebroventricular AAV-GFP and AAV-hCLN7 gene therapy measured at 3-month intervals as histograms above and as a continuous graph below. Analyzed using two-way ANOVA with Tukey post-hoc correction.

**A.** **Rotarod**

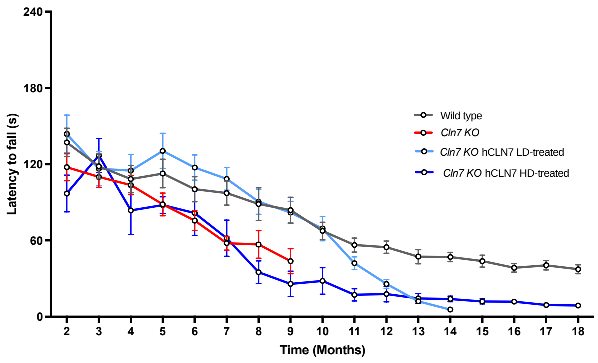


**B.** **Foot Fault**

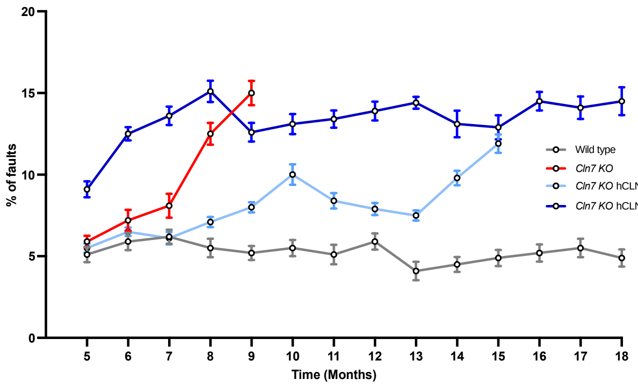


**C.** **Vertical Pole**

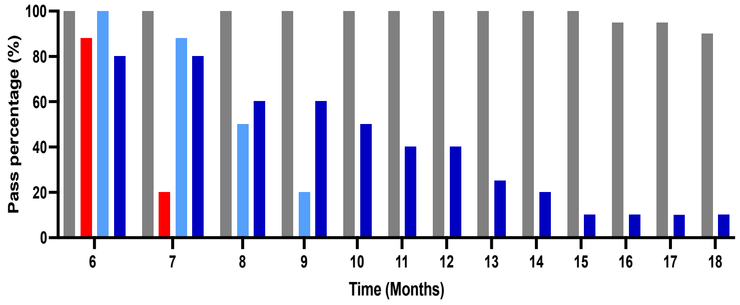


**Fig. S3.** Behavioral analyses for effects of neonatal intracerebroventricular AAV-hCLN7 injections in *Cln7*$^{\Delta ex2}$ mice. **(A)** Accelerating rotarod performance measuring latency to fall, measured at monthly intervals beginning at 2 months. **(B)** Foot-fault measured as percentage of misplaced steps at monthly intervals **(C)** Vertical pole behaviour measured as pass percentage at monthly intervals from 6 months. All data presented as mean ± SEM, (P-values in Supplementary Table 1).

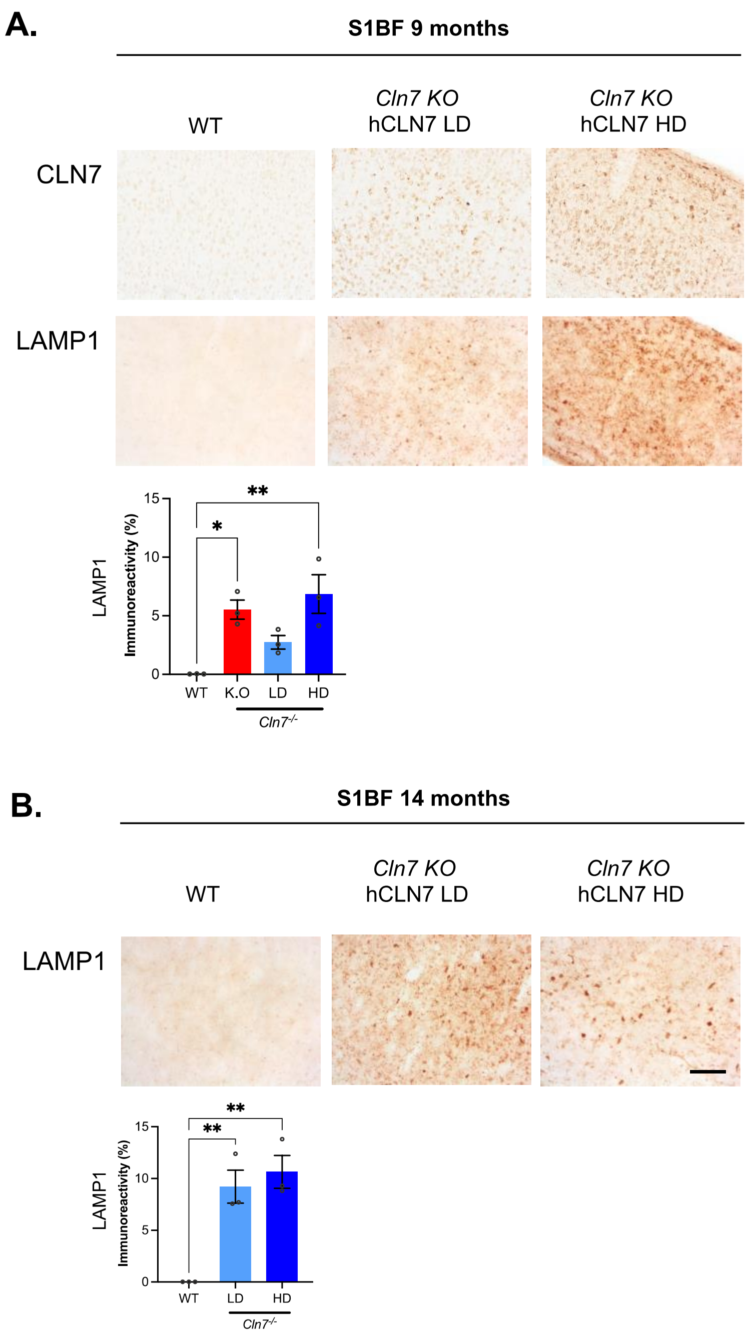


**Fig. S4** Effects of neonatal intracerebroventricular AAV-hCLN7 injections in *Cln7*$^{\Delta ex2}$ mice on CLN7 expression and LAMP1 immunoreactivity. CLN7 and LAMP1 IHC in the S1BF cortex region of wild-type (WT) and *Cln7*$^{\Delta ex2}$ mice receiving low-dose (LD) and High Dose (HD) of AAV-hCLN7 showing dose-dependent increase in expression of CLN7. Representative images at 10x mag and quantified thresholding image analysis at (A) 9 months and (B) 14 months. Scale bar=100µm, analyzed using a one-way ANOVA with post-hoc Bonferroni correction (n=3). All data presented as mean ± SEM, * p≤0.05 and **≤0.01 (P-values in Supplementary Table 1).

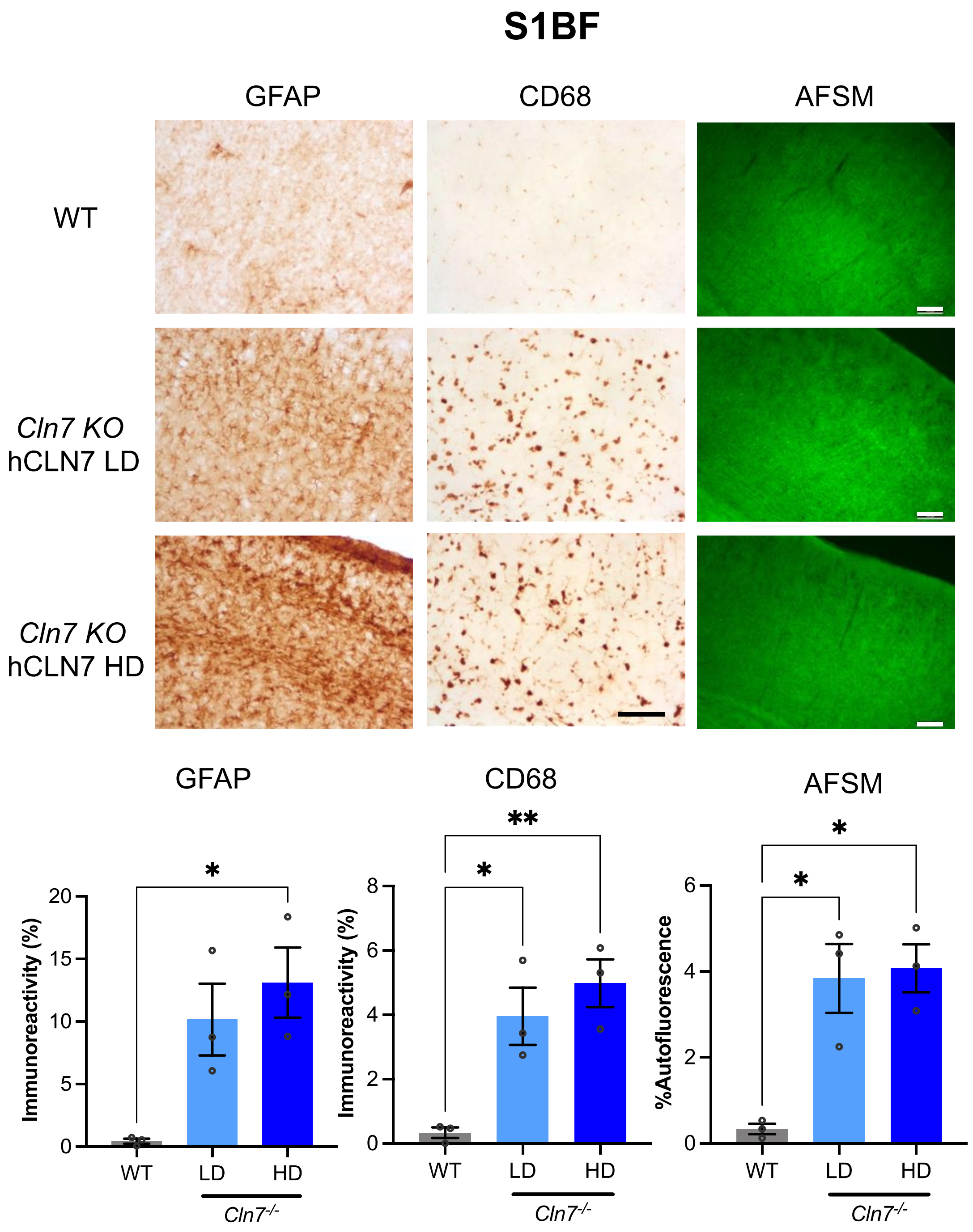


**Fig. S5** Effects of neonatal intracerebroventricular AAV-hCLN7 injections in *Cln7*$^{\Delta ex2}$ mice on neuropathology at 14 months. Representative images and quantified thresholding image analysis at 14 months for GFAP (astrocytes) and CD68 (microglia) immunoreactivity and autofluorescent storage material (AFSM, % autofluorescence) accumulation in the S1BF cortex region. Groups compared are wild-type (WT), low-dose (LD) and high-dose (HD) treated *Cln7*$^{\Delta ex2}$ mice. Scale Bar=100µm, analyzed using a one-way ANOVA with post-hoc Bonferroni correction (n=3). All data presented as mean ± SEM, * p≤0.05 and **≤0.01 (P-values in Supplementary Table 1).

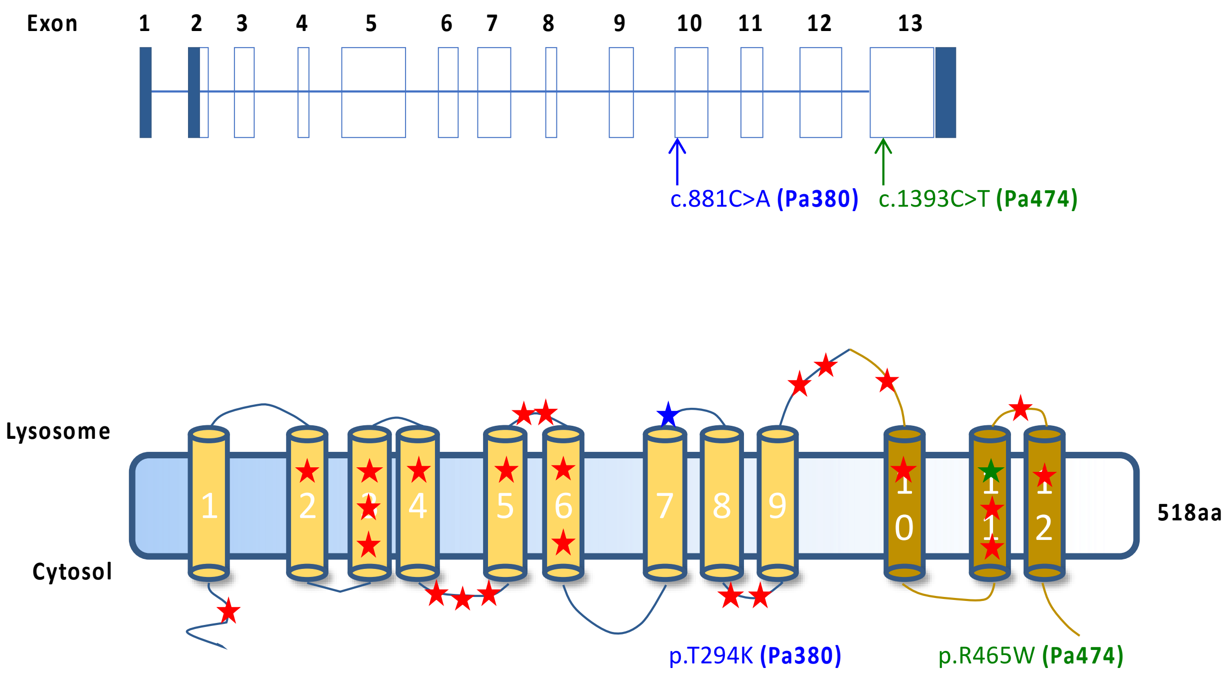


**Fig. S6** Schematic representation of the gene and predicted protein structure for CLN7 including the positions of the homozygous mutations affecting Pa380 and Pa474.

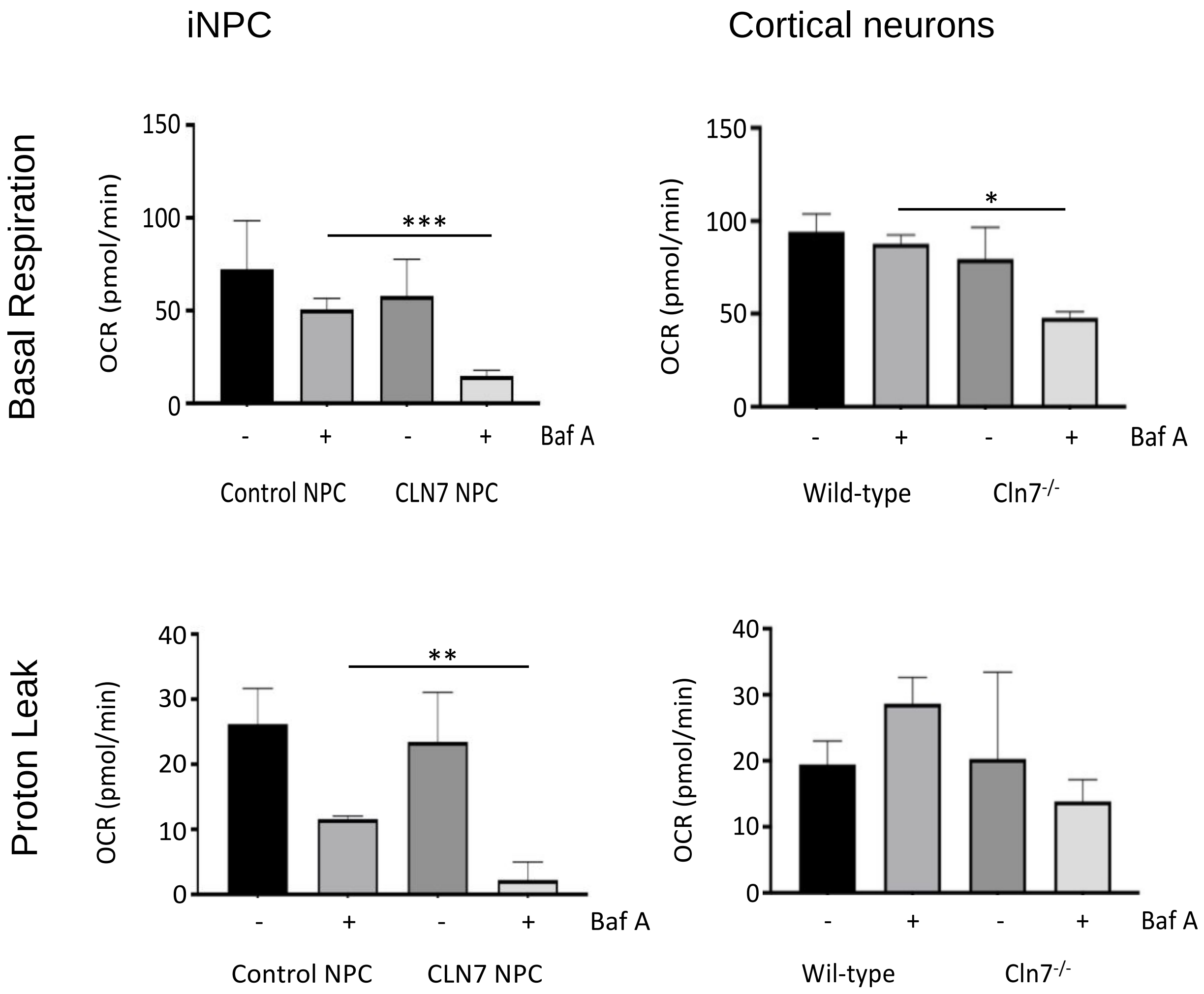


**Fig. S7** Mitochondrial basal respiration and proton leak assayed in the presence or absence of Baf A in CLN7 versus control iNPC (n=6) and *Cln7*$^{\Delta ex2}$ versus wild-type cortical neurons (n=4). Change in oxygen consumption rate (OCR) quantified under steady state conditions for basal respiration using the Seahorse XFp bioanalyer then proton leak was calculated as the difference between basal respiration and ATP-linked respiration after addition of oligomycin. All data presented as mean ± SEM, * p≤0.05, **≤0.01 and ***≤0.001 (P-values in Supplementary Table 1).

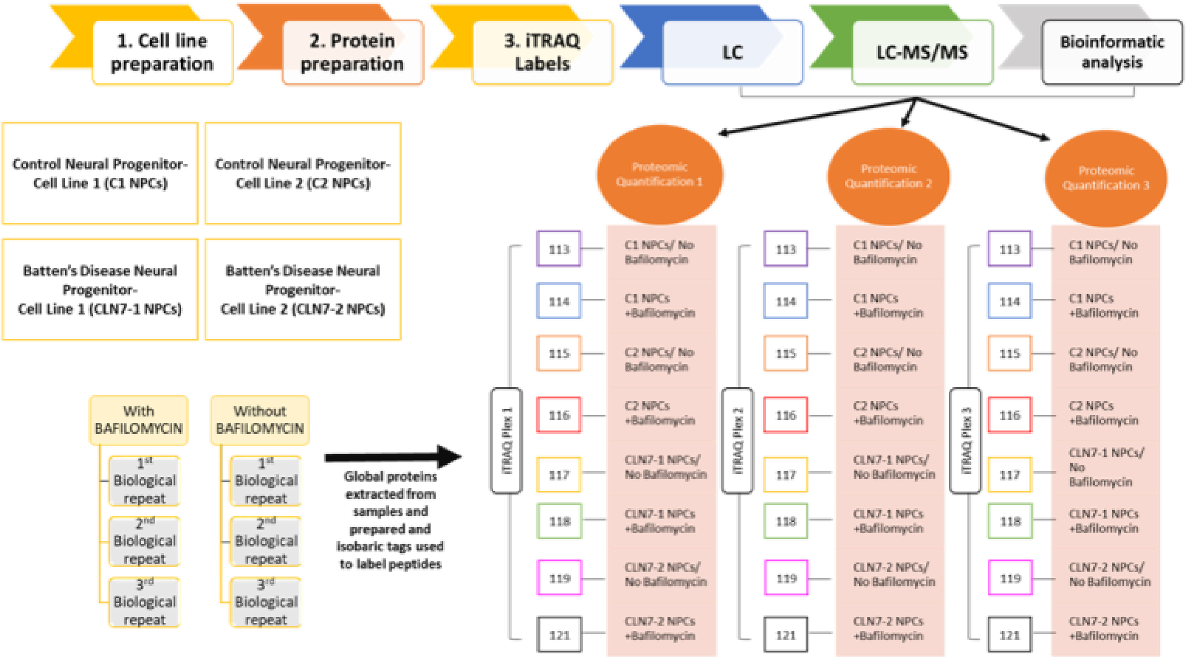


**Fig. S8** Schematic representation of the iTRAQ LC-MS/MS proteomic evaluation.

**A.** Upregulated Basal

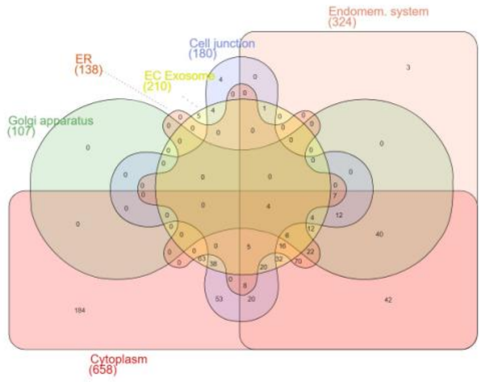


**B.** Downregulated Basal

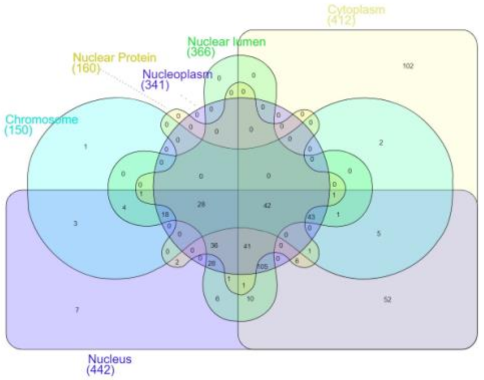


**C.** Upregulated Baf A

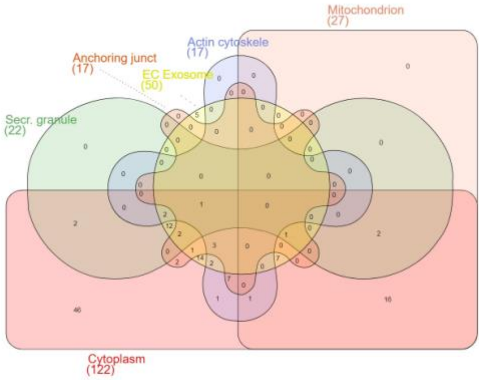


**D.** Downregulated Baf A

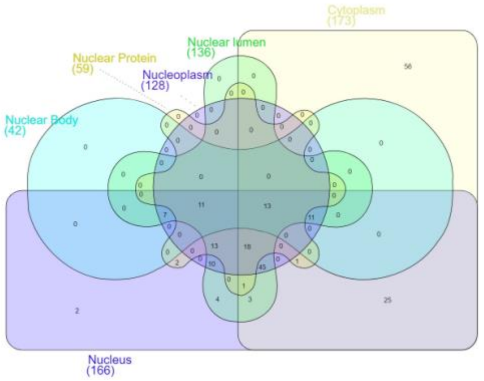


**Fig. S9** Cellular Component Analysis for downregulated proteins under basal condition, upregulated proteins under basal condition, downregulated proteins after Bafilomycin A treatment and upregulated proteins after bafilomycin A treatment.

Upregulated Basal

Downregulated Basal

Upregulated Baf A

Downregulated Baf A

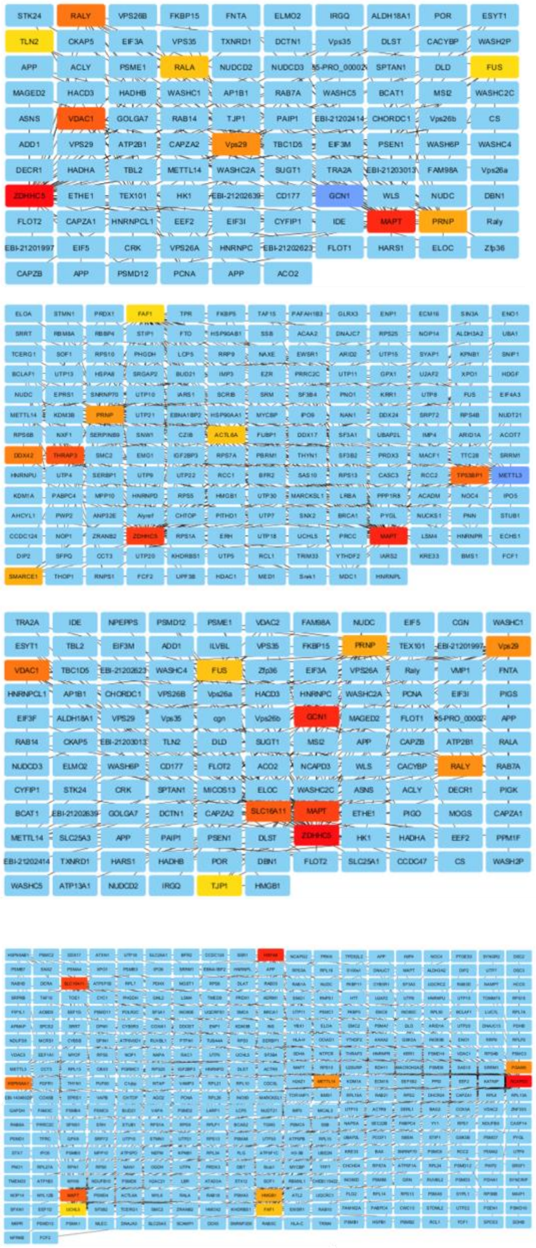


**Fig. S10** Network modelling analysis for downregulated proteins under basal condition, upregulated proteins under basal condition, downregulated proteins after Bafilomycin A treatment and upregulated proteins after bafilomycin A treatment. Network analysis was carried out as described by Stevens *et al* (50). Cytoscape was used to build a network for each list of proteins in the groups mentioned above. Uniprot data base was used to import interactome based on data. The cytoHubba (0.1) Cytoscape Plugin was used to calculate connectivity and bottlenecks for each network. All networks were then ranked separately for both connectivity and bottleneck properties and these were used to evaluate the relative importance of each node and to generate a minimal essential network (MEN). The top 10% of network nodes was chosen to define the MEN, as this represents the most functional elements of the network.

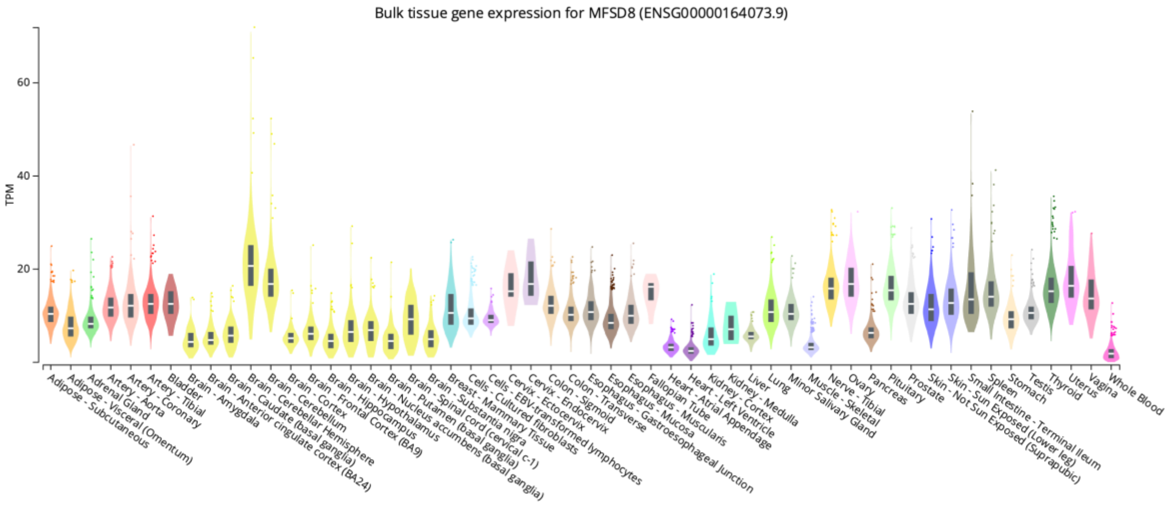


**Fig. S11** Graphical representation of GTEx Portal (gtexportal.org) expression profile for CLN7 transcript (ENSG00000164073.9).

**A.**

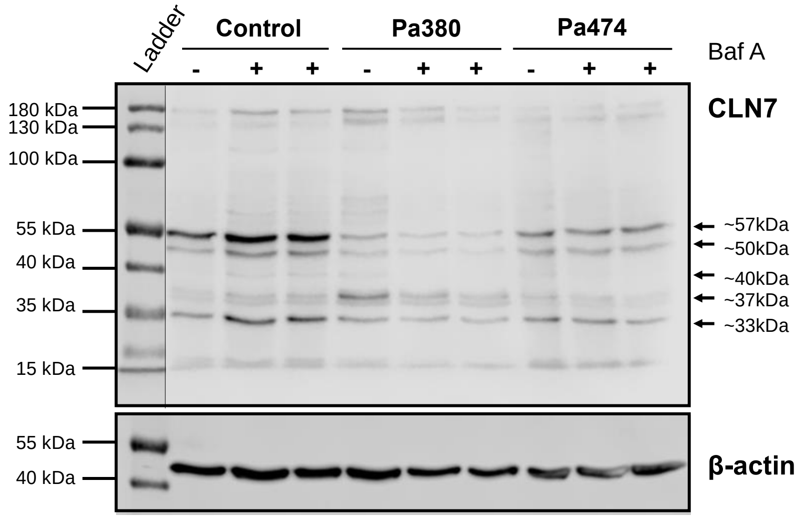


**B.**

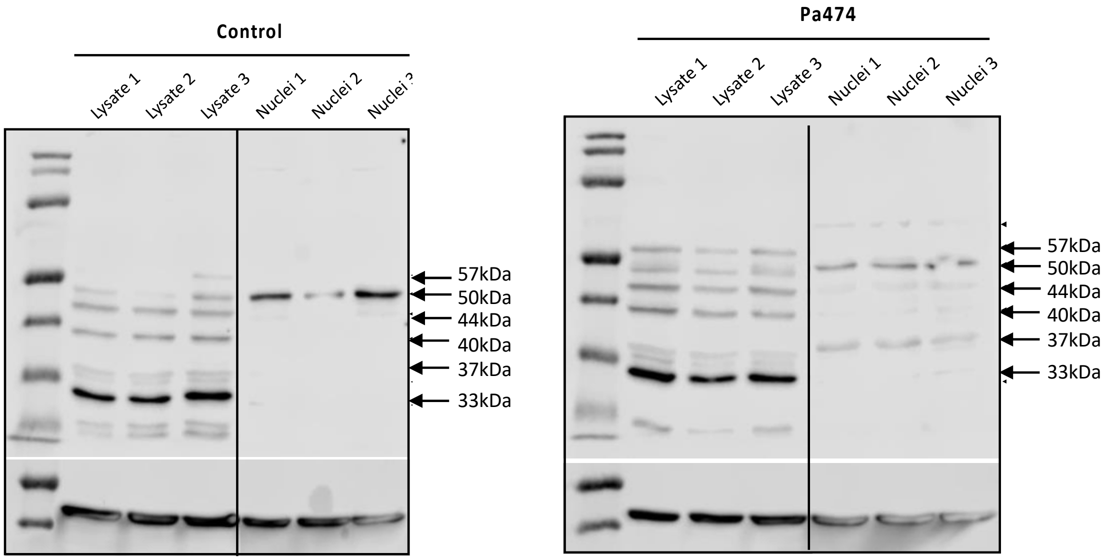


**C.**

| Predicted cleavage site (…) | L | S | K | B | G |
|---|---|---|---|---|---|
| 1 | QAWC†LYTP | AWCL†YTPV | EQAW†CLYT | QAWC†LYTP | AWCL†YTPV |
| 2 | AWCL†YTPV | QAWC†LYTP | AWCL†YTPV | AWCL†YTPV | QAWC†LYTP |
| 3 | SIEQ†AWCL | EQAW†CLYT | QAWC†LYTP | EQAW†CLYT | EQAW†CLYT |

**Fig. S12 (A)** Biological repeats of CLN7 western blots conducted on lysates from CLN7 (Pa380 and Pa474) and control iNPC in the presence and absence of Baf A. (**B**) Biological repeats of CLN7 western blots conducted on whole cell lysates and nuclear fractions from CLN7 (Pa474) and control iNPC. **(C)** Predicted cleavage sites of CLN7 at Cys401 by lysosomal cathepsins L, S, K, B and G.

**Table S1. Experimental p-values**

| Figure | Group | N | Sample Size | Statistical Test | p-value |
|---|---|---|---|---|---|
| 1A | | 82 | WT= 22<br>CLN7 KO= 20<br>CLN7 KO GFP= 11<br>CLN7 KO LD= 18<br>CLN7 KO HD= 11 | Log-rank (Mantel-Cox) test | CLN7 KO vs CLN7 KO GFP = 0.9234, ns<br>CLN7 KO LD vs CLN7 KO HD = 0.1998, ns<br>CLN7 KO vs CLN7 KO LD = <0.0001, ****<br>CLN7 KO vs CLN7 KO HD = 0.0054, **<br>WT vs CLN7 KO= <0.0001, ****<br>WT vs CLN7 KO LD= <0.0054, **<br>WT vs CLN7 KO HD= <0.0767, ns<br>WT vs CLN7 KO GFP= <0.0001, ****<br>CLN7 KO GFP vs CLN7 KO LD = <0.0001, ****<br>CLN7 KO GFP vs CLN7 HD = 0.0136, * |
| 1B | 3-months | 51 | WT=15<br>CLN7 KO= 15<br>CLN7 KO LD= 14<br>CLN7 KO HD= 7 | Two-way ANOVA Tukey's multiple comparison test | WT VS CLN7 KO= 0.9598, ns<br>WT VS CLN7 KO LD= >0.9999, ns<br>WT VS CLN7 KO HD= 0.9165, ns<br>CLN7 KO VS CLN7 KO LD = 0.9757, ns<br>CLN7 KO VS CLN7 KO HD = 0.7023, ns<br>CLN7 KO LD VS CLN7 KO HD = 0.9377, ns |
| 1B | 6-months | 52 | WT= 18<br>CLN7 KO= 13<br>CLN7 KO LD= 14<br>CLN7 KO HD= 7 | Two-way ANOVA Tukey's multiple comparison test | WT VS CLN7 KO= 0.1571, ns<br>WT VS CLN7 KO LD= 0.7486, ns<br>WT VS CLN7 KO HD= 0.7149, ns<br>CLN7 KO VS CLN7 KO LD = 0.0152, *<br>CLN7 KO VS CLN7 KO HD = 0.9894, ns<br>CLN7 KO LD VS CLN7 KO HD = 0.3400, ns |
| 1B | 9-months | 36 | WT= 14<br>CLN7 KO= 7<br>CLN7 KO LD= 9<br>CLN7 KO HD= 6 | Two-way ANOVA Tukey's multiple comparison test | WT VS CLN7 KO= 0.0355, *<br>WT VS CLN7 KO LD= 0.9751, ns<br>WT VS CLN7 KO HD= 0.0036, **<br>CLN7 KO VS CLN7 KO LD = 0.0540, ns<br>CLN7 KO VS CLN7 KO HD = 0.5998, ns<br>CLN7 KO LD VS CLN7 KO HD = 0.0063, ** |
| 1B | 12-months | 26 | WT= 12<br>CLN7 KO LD= 9<br>CLN7 KO HD= 5 | Two-way ANOVA Tukey's multiple comparison test | WT VS CLN7 KO LD= 0.0004, ***<br>WT VS CLN7 KO HD= 0.0023, **<br>CLN7 KO LD VS CLN7 KO HD = 0.5254, ns |
| 1B | 15-months | 18 | WT= 8<br>CLN7 KO LD= 5<br>CLN7 KO HD= 5 | Two-way ANOVA Tukey's multiple comparison test | WT VS CLN7 KO HD= <0.0001, **** |
| 1B | 18-months | 18 | WT= 8<br>CLN7 KO HD= 5 | Two-way ANOVA Tukey's multiple comparison test | WT VS CLN7 KO HD= 0.0002, *** |
| 1C | Foot Fault | 28 | WT= 10<br>CLN7 KO= 8<br>CLN7 KO LD= 5<br>CLN7 KO HD= 5 | One-way ANOVA Tukey´s multiple comparison test | WT VS CLN7 KO= <0.0001, ****<br>WT VS CLN7 KO LD= 0.0591, ns<br>WT VS CLN7 KO HD= <0.0001, ****<br>CLN7 KO VS CLN7 KO LD = 0.0011, **<br>CLN7 KO VS CLN7 KO HD = 0.6307, ns<br>CLN7 KO LD VS CLN7 KO HD = 0.0410, * |
| 1D | Rotations | 23 | WT= 8<br>CLN7 KO= 7<br>CLN7 KO LD= 5<br>CLN7 KO HD= 3 | One-way ANOVA Tukey´s multiple comparison test | WT VS CLN7 KO= 0.9211, ns<br>WT VS CLN7 KO LD= 0.8829, ns<br>WT VS CLN7 KO HD= 0.0274, *<br>CLN7 KO VS CLN7 KO LD = 0.9986, ns<br>CLN7 KO VS CLN7 KO HD = 0.0801, ns<br>CLN7 KO LD VS CLN7 KO HD = 0.1318, ns |

| | | | | | |
|---|---|---|---|---|---|
| **1D** | Time spent in central area | 23 | WT= 8<br>CLN7 KO= 7<br>CLN7 KO LD= 5<br>CLN7 KO HD= 3 | One-way ANOVA Tukey´s multiple comparison test | WT VS CLN7 KO= 0.9713, ns<br>WT VS CLN7 KO LD= 0.9996, ns<br>WT VS CLN7 KO HD= 0.7921, ns<br>CLN7 KO VS CLN7 KO LD = 0.9610, ns<br>CLN7 KO VS CLN7 KO HD = 0.6133, ns<br>CLN7 KO LD VS CLN7 KO HD = 0.8642, ns |
| **1D** | Number of entries in central area | 23 | WT= 8<br>CLN7 KO= 7<br>CLN7 KO LD= 5<br>CLN7 KO HD= 3 | One-way ANOVA Tukey´s multiple comparison test | WT VS CLN7 KO= 0.9691, ns<br>WT VS CLN7 KO LD= 0.4715, ns<br>WT VS CLN7 KO HD= 0.0400, *<br>CLN7 KO VS CLN7 KO LD = 0.7294, ns<br>CLN7 KO VS CLN7 KO HD = 0.0870, ns<br>CLN7 KO LD VS CLN7 KO HD = 0.4250, ns |
| **2A** | Cortical thickness 9-months | 12 | WT= 3<br>CLN7 KO= 3<br>CLN7 KO LD= 3<br>CLN7 KO HD= 3 | One-way ANOVA Bonferroni´s multiple comparison test | WT VS CLN7 KO= 0.0018, **<br>WT VS CLN7 KO LD= 0.1447, ns<br>WT VS CLN7 KO HD= <0.0001, ****<br>CLN7 KO VS CLN7 KO LD = 0.0654, ns<br>CLN7 KO VS CLN7 KO HD = 0.0015, **<br>CLN7 KO LD VS CLN7 KO HD = <0.0001, **** |
| **2A** | Cortical thickness 14-months | 9 | WT= 3<br>CLN7 KO LD= 3<br>CLN7 KO HD= 3 | One-way ANOVA Bonferroni´s multiple comparison test | WT VS CLN7 KO LD= 0.0035, **<br>WT VS CLN7 KO HD= <0.0001, ****<br>CLN7 KO LD VS CLN7 KO HD = 0.0018, ** |
| **2D** | ASFM in S1BF | 12 | WT= 3<br>CLN7 KO= 3<br>CLN7 KO LD= 3<br>CLN7 KO HD= 3 | One-way ANOVA Bonferroni´s multiple comparison test | WT VS CLN7 KO= 0.0078, **<br>WT VS CLN7 KO LD= 0.2203, ns<br>WT VS CLN7 KO HD= 0.2308, ns<br>CLN7 KO VS CLN7 KO LD = 0.2901, ns<br>CLN7 KO VS CLN7 KO HD = 0.2769, ns<br>CLN7 KO LD VS CLN7 KO HD = >0.9999, ns |
| **2D** | GFAP in S1BF | 12 | WT= 3<br>CLN7 KO= 3<br>CLN7 KO LD= 3<br>CLN7 KO HD= 3 | One-way ANOVA Bonferroni´s multiple comparison test | WT VS CLN7 KO= <0.0001, ****<br>WT VS CLN7 KO LD= 0.0079, **<br>WT VS CLN7 KO HD= <0.0001, ****<br>CLN7 KO VS CLN7 KO LD = 0.0048, **<br>CLN7 KO VS CLN7 KO HD = >0.9999, ns<br>CLN7 KO LD VS CLN7 KO HD = 0.0012, ** |
| **2D** | CD68 in S1BF | 12 | WT= 3<br>CLN7 KO= 3<br>CLN7 KO LD= 3<br>CLN7 KO HD= 3 | One-way ANOVA Bonferroni´s multiple comparison test | WT VS CLN7 KO= 0.0008, ***<br>WT VS CLN7 KO LD= >0.9999, ns<br>WT VS CLN7 KO HD= 0.0238, *<br>CLN7 KO VS CLN7 KO LD = 0.0032, **<br>CLN7 KO VS CLN7 KO HD = 0.1406, ns<br>CLN7 KO LD VS CLN7 KO HD = 0.1433, ns |
| **4A** | Mitochondrial Membrane Potential | 8 | Cont NPC=6<br>CLN7 NPC=6 | Two-tailed t-test | Cont NPC vs CLN7 NPC=0.0036, ** |
| **4B** | MitoSOX | 8 | Cont NPC=6<br>Cont NPC + Baf=6<br>CLN7 NPC=6<br>CLN7 NPC + Baf=6 | Two-tailed t-test | Cont NPC vs CLN7 NPC=0.0017, ** |
| **4C** | MitoStress Test | 6/4 | Cont NPC=6<br>Cont NPC + Baf=6<br>CLN7 NPC=6<br>CLN7 NPC + Baf=6<br>WT=4<br>WT + Baf=4<br>CLN7 KO=4<br>CLN7 KO + Baf=4 | Two-tailed t-test | <u>ATP Production</u><br>Cont NPC +Baf vs CLN7 NPC +Baf=0.0012, **<br>WT +Baf vs CLN7 KO +Baf=0.0452, *<br><u>Maximal Respiration</u><br>Cont NPC +Baf vs CLN7 NPC +Baf=0.0066, **<br>WT +Baf vs CLN7 KO +Baf=0.0450, * |
| **4D** | MTT Assay | 6 | Cont NPC=6<br>Cont NPC + Baf=6 | One-tailed t-test | CLN7 NPC vs CLN7 NPC +Baf=0.0001<br>Cont NPC +Baf vs CLN7 NPC +Baf=0.0311 |

| | | | | | |
|---|---|---|---|---|---|
| | | | CLN7 NPC=6<br>CLN7 NPC + Baf=6 | | |
| **4E** | qRT-PCR | 4 | Cont NPC=3<br>CLN7 NPC=3 | One-tailed t-test | ATP5A<br>Cont NPC vs CLN7 NPC=0.0003, ***<br>UCP2<br>Cont NPC vs CLN7 NPC=0.0473, * |
| **5C** | Apotracker | 3 | Cont NPC=3 (3)<br>Cont NPC + Baf=3 (3)<br>Pa380 CLN7 NPC=3 (3)<br>Pa380 CLN7 NPC + Baf=3 (3)<br>Pa474 CLN7 NPC=3 (3)<br>Pa474 CLN7 NPC + Baf=3 (3) | One-tailed t-test | Cont NPC vs Cont NPC +Baf= n.s.<br>Pa380 NPC vs Pa380 NPC +Baf=$1.216x10^{-5}$, ****<br>Pa474 NPC vs Pa474 NPC +Baf=$1.684x10^{-5}$, **** |
| **8E** | Apotracker | 3 | HEK293T =3<br>HEK293T + C1 =3<br>HEK293T + e7/8 =3<br>HEK293T + e7/8C =3 | One-tailed t-test | HEK293T vs HEK293T<br>HEK293T vs HEK293T + C1<br>HEK293T vs HEK293T + e7/8<br>HEK293T vs HEK293T + e7/8c1 |
| **S2** | 3-months | 89 | WT= 22<br>CLN7 KO= 27<br>CLN7 KO GFP= 11<br>CLN7 KO LD= 18<br>CLN7 KO HD= 11 | Two-way ANOVA Tukey´s multiple comparison test | WT vs CLN7 KO= 0.9993, ns<br>WT vs CLN7 KO GFP= 0.9999, ns<br>WT vs CLN7 KO LD= 0.9839, ns<br>WT vs CLN7 KO HD= 0.5719, ns<br>CLN7 KO vs CLN7 KO GFP= 0.9973, ns<br>CLN7 KO vs CLN7 KO LD= 0.9978, ns<br>CLN7 KO vs CLN7 KO HD= 0.4731, ns<br>CLN7 KO GFP vs CLN7 KO LD= 0.9796, ns<br>CLN7 KO GFP vs CLN7 KO HD= 0.7809, ns<br>CLN7 KO LD vs CLN7 KO HD= 0.3870, ns |
| **S2** | 6-months | 74 | WT= 20<br>CLN7 KO= 18<br>CLN7 KO GFP= 9<br>CLN7 KO LD= 17<br>CLN7 KO HD= 10 | Two-way ANOVA Tukey´s multiple comparison test | WT vs CLN7 KO= >0.9999, ns<br>WT vs CLN7 KO GFP= 0.8464, ns<br>WT vs CLN7 KO LD= 0.9776, ns<br>WT vs CLN7 KO HD= 0.6891, ns<br>CLN7 KO vs CLN7 KO GFP= 0.8861, ns<br>CLN7 KO vs CLN7 KO LD= 0.9839, ns<br>CLN7 KO vs CLN7 KO HD= 0.7511, ns<br>CLN7 KO GFP vs CLN7 KO LD= 0.6127, ns<br>CLN7 KO GFP vs CLN7 KO HD= 0.9978, ns<br>CLN7 KO LD vs CLN7 KO HD= 0.4565, ns |
| **S2** | 9-months | 44 | WT= 13<br>CLN7 KO= 4<br>CLN7 KO GFP= 7<br>CLN7 KO LD= 13<br>CLN7 KO HD= 7 | Two-way ANOVA Tukey´s multiple comparison test | WT vs CLN7 KO= 0.3264, ns<br>WT vs CLN7 KO GFP= 0.0845, ns<br>WT vs CLN7 KO LD= 0.9384, ns<br>WT vs CLN7 KO HD= 0.3342, ns<br>CLN7 KO vs CLN7 KO GFP= 0.9978, ns<br>CLN7 KO vs CLN7 KO LD= 0.5443, ns<br>CLN7 KO vs CLN7 KO HD= 0.9071, ns<br>CLN7 KO GFP vs CLN7 KO LD= 0.3268, ns<br>CLN7 KO GFP vs CLN7 KO HD= 0.9212, ns<br>CLN7 KO LD vs CLN7 KO HD= 0.7793, ns |
| **S2** | 12-months | 26 | WT= 12<br>CLN7 KO LD= 13<br>CLN7 KO HD= 5 | Two-way ANOVA Tukey´s multiple comparison test | WT vs CLN7 KO LD= 0.0725, ns<br>WT vs CLN7 KO HD= 0.1243, ns<br>CLN7 KO LD vs CLN7 KO HD= 0.9540, ns |
| **S2** | 15-months | 18 | WT=4<br>CLN7 KO HD= 4 | Two-way ANOVA Tukey´s multiple comparison test | WT vs CLN7 KO HD= 0.1859, ns |

| | | | | | |
|---|---|---|---|---|---|
| **S2** | 18-months | 18 | WT=2<br>CLN7 KO HD= 3 | Two-way ANOVA<br>Tukey´s multiple comparison test | WT vs CLN7 KO HD= 0.4354, ns |
| **S3** | Time mobile | 23 | WT= 8<br>CLN7 KO= 7<br>CLN7 KO LD= 5<br>CLN7 KO HD= 3 | One-way ANOVA<br>Tukey´s multiple comparison test | WT VS CLN7 KO= 0.3941, ns<br>WT VS CLN7 KO LD= 0.4097, ns<br>WT VS CLN7 KO HD= 0.1382, ns<br>CLN7 KO VS CLN7 KO LD = 0.9994, ns<br>CLN7 KO VS CLN7 KO HD = 0.7370, ns<br>CLN7 KO LD VS CLN7 KO HD = 0.8174, ns |
| **S3** | Distance travelled | 23 | WT= 8<br>CLN7 KO= 7<br>CLN7 KO LD= 5<br>CLN7 KO HD= 3 | One-way ANOVA<br>Tukey´s multiple comparison test | WT VS CLN7 KO= 0.8632, ns<br>WT VS CLN7 KO LD= 0.8222, ns<br>WT VS CLN7 KO HD= 0.0386, *<br>CLN7 KO VS CLN7 KO LD = 0.9986, ns<br>CLN7 KO VS CLN7 KO HD = 0.1333, ns<br>CLN7 KO LD VS CLN7 KO HD = 0.2056, ns |
| **S3** | Speed | 23 | WT= 8<br>CLN7 KO= 7<br>CLN7 KO LD= 5<br>CLN7 KO HD= 3 | One-way ANOVA<br>Tukey´s multiple comparison test | WT VS CLN7 KO= 0.8694, ns<br>WT VS CLN7 KO LD= 0.8273, ns<br>WT VS CLN7 KO HD= 0.0393, *<br>CLN7 KO VS CLN7 KO LD = 0.9986, ns<br>CLN7 KO VS CLN7 KO HD = 0.1328, ns<br>CLN7 KO LD VS CLN7 KO HD = 0.2057, ns |
| **S4** | LAMP1 in S1BF | 12 | WT= 3<br>CLN7 KO= 3<br>CLN7 KO LD= 3<br>CLN7 KO HD= 3 | One-way ANOVA<br>Bonferroni´s multiple comparison test | WT VS CLN7 KO= 0.0232, *<br>WT VS CLN7 KO LD= 0.5011, ns<br>WT VS CLN7 KO HD= 0.0064, **<br>CLN7 KO VS CLN7 KO LD = 0.4556, ns<br>CLN7 KO VS CLN7 KO HD = >0.9999, ns<br>CLN7 KO LD VS CLN7 KO HD = 0.1001, ns |
| **S4** | LAMP1 in S1BF | 9 | WT= 3<br>CLN7 KO LD= 3<br>CLN7 KO HD= 3 | One-way ANOVA<br>Bonferroni´s multiple comparison test | WT VS CLN7 KO LD= 0.0072, **<br>WT VS CLN7 KO HD= 0.0035, **<br>CLN7 KO LD VS CLN7 KO HD = >0.9999, ns |
| **S5** | GFAP in S1BF | 9 | WT= 3<br>CLN7 KO LD= 3<br>CLN7 KO HD= 3 | One-way ANOVA<br>Bonferroni´s multiple comparison test | WT VS CLN7 KO LD= 0.0752, ns<br>WT VS CLN7 KO HD= 0.0248, *<br>CLN7 KO LD VS CLN7 KO HD = >0.9999, ns |
| **S5** | CD68 in S1BF | 9 | WT= 3<br>CLN7 KO LD= 3<br>CLN7 KO HD= 3 | One-way ANOVA<br>Bonferroni´s multiple comparison test | WT VS CLN7 KO LD= 0.0274, *<br>WT VS CLN7 KO HD= 0.0085, **<br>CLN7 KO LD VS CLN7 KO HD = 0.9717, ns |
| **S5** | AFSM in S1BF | 9 | WT= 3<br>CLN7 KO LD= 3<br>CLN7 KO HD= 3 | One-way ANOVA<br>Bonferroni´s multiple comparison test | WT VS CLN7 KO LD= 0.0144, *<br>WT VS CLN7 KO HD= 0.0105, *<br>CLN7 KO LD VS CLN7 KO HD = >0.9999, ns |
| **S7** | MitoStress Test | 6<br>(4) | Cont NPC=6<br>Cont NPC + Baf=6<br>CLN7 NPC=6<br>CLN7 NPC + Baf=6<br>WT=4<br>WT + Baf=4<br>CLN7 KO=4<br>CLN7 KO + Baf=4 | Two-tailed t-test | <u>Basal Respiration</u><br>Cont NPC +Baf vs CLN7 NPC +Baf=0.0007, ***<br>WT +Baf vs CLN7 KO +Baf=0.013, *<br><u>Proton Leak</u><br>Cont NPC +Baf vs CLN7 NPC +Baf=0.0045, **<br>WT +Baf vs CLN7 KO +Baf=0.0043, ** |

### Table S2. Plasmid expression vector sequences

| Plasmid | | Sequence |
|---|---|---|
| **pcDNA3.4-cMFSD8** | DNA | ATGGCCGGCCTGAGAAACGAGTCTGAGCAAGAGCCTCTGCTGGGCGATACACCTGGCAGCAGAGAGTGGGACATCCTGGAAACCGAGGAACACT<br>ACAAGAGCCGGTGGCGGAGCATCCGGATCCTGTACCTGACCATGTTCCTGAGCAGCGTGGGCTTCAGCGTGGTCATGATGAGCATCTGGCCCTACC<br>TGCAGAAGATCGACCCTACCGCCGATACCAGCTTCCTCGGATGGGTTATCGCCAGCTACAGCCTGGGACAGATGGTGGCCTCTCCTATCTTTGGCCT<br>GTGGTCCAACTACAGACCCCGGAAAGAGCCCCTGATCGTCAGCATCCTGATTAGCGTGGCCGCCAACTGCCTGTACGCCTATCTGCACATTCCCGCC<br>AGCCACAACAAGTACTACATGCTGGTGGCCAGAGGCCTGCTCGGAATCGGAGCTGGAAATGTGGCCGTCGTGCGGAGCTATACAGCCGGCGCTAC<br>AAGCCTGCAAGAGCGGACAAGCAGCATGGCCAACATCAGCATGTGTCAGGCCCTGGGCTTCATTCTGGGCCCTGTGTTCCAGACCTGCTTCACCTTT<br>CTGGGCGAGAAGGGCGTGACCTGGGACGTGATCAAGCTGCAGATCAACATGTACACCACACCTGTGCTGCTGAGCGCCTTCCTGGGCATCCTGAAC<br>ATCATCCTGATCCTGGCCATTCTGCGCGAGCACAGAGTGGACGATTCTGGCAGACAGTGCAAGAGCATCAACTTCGAGGAAGCCAGCACCGACGA<br>GGCTCAGGTGCCACAGGGAAACATTGATCAGGTGGCCGTGGTGGCCATCAACGTGCTGTTCTTCGTGACCCTGTTCATCTTCGCCCTGTTCGAGACA<br>ATCATCACCCCTCTGACCATGGATATGTACGCCTGGACACAAGAGCAGGCCGTGCTGTACAACGGAATCATCCTGGCCGCTCTGGGCGTTGAGGCC<br>GTGGTTATTTTCCTGGGCGTGAAGCTGCTGTCCAAGAAGATCGGCGAGAGAGCCATCCTGCTCGGCGGACTGATCGTTGTGTGGGTCGGATTCTTC<br>ATCCTGCTGCCTTGGGGCAATCAGTTCCCCAAGATCCAGTGGGAAGATCTGCACAACAACAGCATCCCCAACACCACCTTCGGCGAGATCATCATCG<br>GCCTGTGGAAGTCCCCAATGGAAGATGACAACGAGCGGCCCACAGGCTGA |
| | Protein | MAGLRNESEQEPLLGDTPGSREWDILETEEHYKSRWRSIRILYLTMFLSSVGFSVVMMSIWPYLQKIDPTADTSFLGWVIASYSLGQMVASPIFGLWSNYR<br>PRKEPLIVSILISVAANCLYAYLHIPASHNKYYMLVARGLLGIGAGNVAVVRSYTAGATSLQERTSSMANISMCQALGFILGPVFQTCFTFLGEKGVTWDVIK<br>LQINMYTTPVLLSAFLGILNIILILAILREHRVDDSGRQCKSINFEEASTDEAQVPQGNIDQVAVVAINVLFFVTLFIFALFETIITPLTMDMYAWTQEQAVLYN<br>GIILAALGVEAVVIFLGVKLLSKKIGERAILLGGLIVVWVGFFILLPWGNQFPKIQWEDLHNNSIPNTTFGEIIIGLWKSPMEDDNERPTG* |
| **pcDNA3.4-e78MFSD8** | DNA | ATGGCCGGCCTGAGAAACGAGTCTGAGCAAGAGCCTCTGCTGGGCGATACACCTGGCAGCAGAGAGTGGGACATCCTGGAAACCGAGGAACACT<br>ACAAGAGCCGGTGGCGGAGCATCCGGATCCTGTACCTGACCATGTTCCTGAGCAGCGTGGGCTTCAGCGTGGTCATGATGAGCATCTGGCCCTACC<br>TGCAGAAGATCGACCCTACCGCCGATACCAGCTTCCTCGGATGGGTTATCGCCAGCTACAGCCTGGGACAGATGGTGGCCTCTCCTATCTTTGGCCT<br>GTGGTCCAACTACAGACCCCGGAAAGAGCCCCTGATCGTCAGCATCCTGATTAGCGTGGCCGCCAACTGCCTGTACGCCTATCTGCACATTCCCGCC<br>AGCCACAACAAGTACTACATGCTGGTGGCCAGAGGCCTGCTCGGAATCGGAGCTGGAAATGTGGCCGTCGTGCGGAGCTATACAGCCGGCGCTAC<br>AAGCCTGCAAGAGCGGACAAGCAGCATGGCCAACATCAGCATGTGTCAGGCCCTGGGCTTCATCCTGGGACCTGCCTCTACAGATGAGGCCCAGG<br>TGCCACAGGGCAACATTGATCAAGTGGCCGTGGTGGCCATCAACGTGCTGTTCTTCGTGACCCTGTTCATCTTCGCCCTGTTCGAGACAATCATCACC<br>CCTCTGACCATGGATATGTACGCCTGGACACAAGAGCAGGCCGTGCTGTACAACGGCATCATTCTGGCCGCTCTGGGCGTTGAGGCCGTGGTTATT<br>TTTCTGGGCGTGAAGCTGCTGTCCAAGAAGATCGGCGAGAGAGCCATTCTGCTCGGCGGCCTGATCGTTGTGTGGGTCGGATTCTTCATTCTGCTGC<br>CCTGGGGCAATCAGTTCCCCAAGATCCAGTGGGAAGATCTGCACAACAACAGCATCCCCAACACCACCTTCGGCGAGATCATCATCGGCCTGTGGA<br>AGTCCCCAATGGAAGATGACAACGAGAGGCCCACCGGCTGCTCTATTGAACAGGCCTGGTGTCTGTACACCCCTGTGATCCACCTGGCTCAGTTTCT<br>GACAAGCGCCGTGCTGATCGGCCTGGGCTACCCTGTGTGTAACCTGATGAGCTACACCCTGTACAGCAAGATTCTGGGCCCCAAGCCTCAGGGCGT<br>GTACATGGGATGGCTGACAGCTTCTGGAAGCGGCGCCAGAATCCTGGGGCCTATGTTTATCAGTCAGGTGTACGCTCACTGGGGCCCTAGATGGGC<br>CTTTTCTCTCGTGTGCGGCATCATCGTGCTGACCATCACACTGCTGGGAGTCGTGTACAAGCGGCTGATCGCCCTGAGCGTCAGATACGGCAGAATC<br>CAAGAGTGA |
| | Protein | MAGLRNESEQEPLLGDTPGSREWDILETEEHYKSRWRSIRILYLTMFLSSVGFSVVMMSIWPYLQKIDPTADTSFLGWVIASYSLGQMVASPIFGLWSNYR<br>PRKEPLIVSILISVAANCLYAYLHIPASHNKYYMLVARGLLGIGAGNVAVVRSYTAGATSLQERTSSMANISMCQALGFILGPASTDEAQVPQGNIDQVAVV<br>AINVLFFVTLFIFALFETIITPLTMDMYAWTQEQAVLYNGIILAALGVEAVVIFLGVKLLSKKIGERAILLGGLIVVWVGFFILLPWGNQFPKIQWEDLHNNSIP<br>NTTFGEIIIGLWKSPMEDDNERPTGCSIEQAWCLYTPVIHLAQFLTSAVLIGLGYPVCNLMSYTLYSKILGPKPQGVYMGWLTASGSGARILGPMFISQVYA<br>HWGPRWAFSLVCGIIVLTITLLGVVYKRLIALSVRYGRIQE* |
| **pcDNA3.4-e78MFSD8c** | DNA | ATGGCCGGCCTGAGAAACGAGTCTGAGCAAGAGCCTCTGCTGGGCGATACACCTGGCAGCAGAGAGTGGGACATCCTGGAAACCGAGGAACACT<br>ACAAGAGCCGGTGGCGGAGCATCCGGATCCTGTACCTGACCATGTTCCTGAGCAGCGTGGGCTTCAGCGTGGTCATGATGAGCATCTGGCCCTACC<br>TGCAGAAGATCGACCCTACCGCCGATACCAGCTTCCTCGGATGGGTTATCGCCAGCTACAGCCTGGGACAGATGGTGGCCTCTCCTATCTTTGGCCT<br>GTGGTCCAACTACAGACCCCGGAAAGAGCCCCTGATCGTCAGCATCCTGATTAGCGTGGCCGCCAACTGCCTGTACGCCTATCTGCACATTCCCGCC<br>AGCCACAACAAGTACTACATGCTGGTGGCCAGAGGCCTGCTCGGAATCGGAGCTGGAAATGTGGCCGTCGTGCGGAGCTATACAGCCGGCGCTAC<br>AAGCCTGCAAGAGCGGACAAGCAGCATGGCCAACATCAGCATGTGTCAGGCCCTGGGCTTCATCCTGGGACCTGCCTCTACAGATGAGGCCCAGG<br>TGCCACAGGGCAACATTGATCAAGTGGCCGTGGTGGCCATCAACGTGCTGTTCTTCGTGACCCTGTTCATCTTCGCCCTGTTCGAGACAATCATCACC<br>CCTCTGACCATGGATATGTACGCCTGGACACAAGAGCAGGCCGTGCTGTACAACGGCATCATTCTGGCCGCTCTGGGCGTTGAGGCCGTGGTTATT<br>TTTCTGGGCGTGAAGCTGCTGTCCAAGAAGATCGGCGAGAGAGCCATTCTGCTCGGCGGCCTGATCGTTGTGTGGGTCGGATTCTTCATTCTGCTGC<br>CCTGGGGCAATCAGTTCCCCAAGATCCAGTGGGAAGATCTGCACAACAACAGCATCCCCAACACCACCTTCGGCGAGATCATCATCGGCCTGTGGA<br>AGTCCCCAATGGAAGATGACAACGAGCGGCCCACAGGCTGA |
| | Protein | MAGLRNESEQEPLLGDTPGSREWDILETEEHYKSRWRSIRILYLTMFLSSVGFSVVMMSIWPYLQKIDPTADTSFLGWVIASYSLGQMVASPIFGLWSNYR<br>PRKEPLIVSILISVAANCLYAYLHIPASHNKYYMLVARGLLGIGAGNVAVVRSYTAGATSLQERTSSMANISMCQALGFILGPASTDEAQVPQGNIDQVAVV<br>AINVLFFVTLFIFALFETIITPLTMDMYAWTQEQAVLYNGIILAALGVEAVVIFLGVKLLSKKIGERAILLGGLIVVWVGFFILLPWGNQFPKIQWEDLHNNSIP<br>NTTFGEIIIGLWKSPMEDDNERPTG* |
